# A long-deletion mouse model of Williams syndrome reveals *Ncf1*-dependent modulation of vascular and neural phenotypes

**DOI:** 10.1101/2023.10.30.564727

**Authors:** Laurens W.J. Bosman, Hamid el Azzouzi, Lieke Kros, Nicole van Vliet, Yanto Ridwan, Stéphanie Dijkhuizen, Erika Goedknegt, Bastian S. Generowicz, Martijn C. Sierksma, Dick Jaarsma, Manuele Novello, Morrisen Snoeren, Emma Kretschmann, Danique Broere, Rocco Caliandro, Sebastiaan K.E. Koekkoek, Pieter Kruizinga, Vera van Dis, Haibo Zhou, Hui Yang, Changyang Zhou, Ingrid van der Pluijm, Jeroen Essers, Chris I. De Zeeuw

## Abstract

Williams syndrome is a developmental disorder caused by a microdeletion entailing the loss of a single copy of 25-27 genes on chromosome 7q11.23. Patients suffer from cardiovascular and neuropsychological symptoms. Structural abnormalities of the cardiovascular system in Williams syndrome have been attributed to the hemizygous loss of the elastin (*ELN*) gene. In contrast, the neuropsychological consequences of Williams syndrome, including sensorimotor deficits, hypersociability and cognitive impairments, have been mainly attributed to altered expression of transcription factors like *LIMK1*, *GTF2I* and *GTF2IRD1*, while the potential impact of altered cerebrovascular function has been largely overlooked. To study the relationship between Williams syndrome mutations and vascularization of both the heart and brain, we generated a mouse model carrying a relatively long microdeletion (LD) that includes the *Ncf1* gene, thereby minimizing the confounding impact of hypertension. LD mice had elongated and tortuous aortas but, unlike *Eln* haploinsufficient mice, showed no signs of structural cardiac hypertrophy. Remarkably, LD mice also displayed structural abnormalities in coronary and brain vessels, including disorganized extracellular matrices. Importantly, LD mice faithfully replicated both cardiovascular and neuropsychological symptoms observed in patients. The phenotype was even more comprehensive than former models, with structure-function correlations evident in aberrant auditory and motor behaviors resembling those in patients with Williams syndrome. Together, our findings suggest that not only cardiovascular but also neuropsychological symptoms in Williams syndrome may be driven in part by vascular abnormalities affecting both heart and brain.

**Significance Statement:** Williams syndrome is caused by microdeletion of 25-27 genes on chromosome 7q11.23, resulting in cardiovascular and neuropsychological symptoms. It remains unclear how the affected genes interact and whether cardiovascular deficits influence brain function. We developed and characterized a mouse model with the longest Williams syndrome microdeletion to date. This model reveals interactions between genes that can be compensatory or additive: haploinsufficiency of *Ncf1* may counteract the cardiac hypertrophy caused by *Eln* deletion, while vascular defects that are potentially due to *Eln* haploinsufficiency extend to the brain and may worsen neuropsychological symptoms. Our findings support the hypothesis that structural vascular deficits putatively contribute to both cardiac and cognitive phenotypes in Williams syndrome, opening new avenues for understanding and treating this syndrome.

## Introduction

Williams syndrome, also known as Williams-Beuren syndrome, is a developmental multisystem disorder caused by a hemizygous microdeletion in the Williams syndrome critical region (WSCR) on chromosome 7q11.23 (Peoples et al., 2000; Bayés et al., 2003; Kozel et al., 2021). Most patients have a microdeletion involving 25 genes (1.55 Mb), some have a longer deletion encompassing 27 genes (1.85 Mb), while occasionally atypical shorter or longer deletions occur (Peoples et al., 2000; Bayés et al., 2003; Fusco et al., 2014; Lugo et al., 2020; Kozel et al., 2021) (Fig. 1A). Patients with Williams syndrome exhibit a wide range of symptoms, including cardiovascular abnormalities, distinctive facial features, hypersocial behavior, mild to moderate intellectual disability, hyperacusis and mild motor deficits (Williams et al., 1961; Beuren et al., 1962; Klein et al., 1990; Hoogenraad et al., 2004; Gothelf et al., 2006; van Hagen et al., 2007; Collins et al., 2010; Kozel et al., 2021; Vernerova and Solc, 2024).

**Figure 1.**
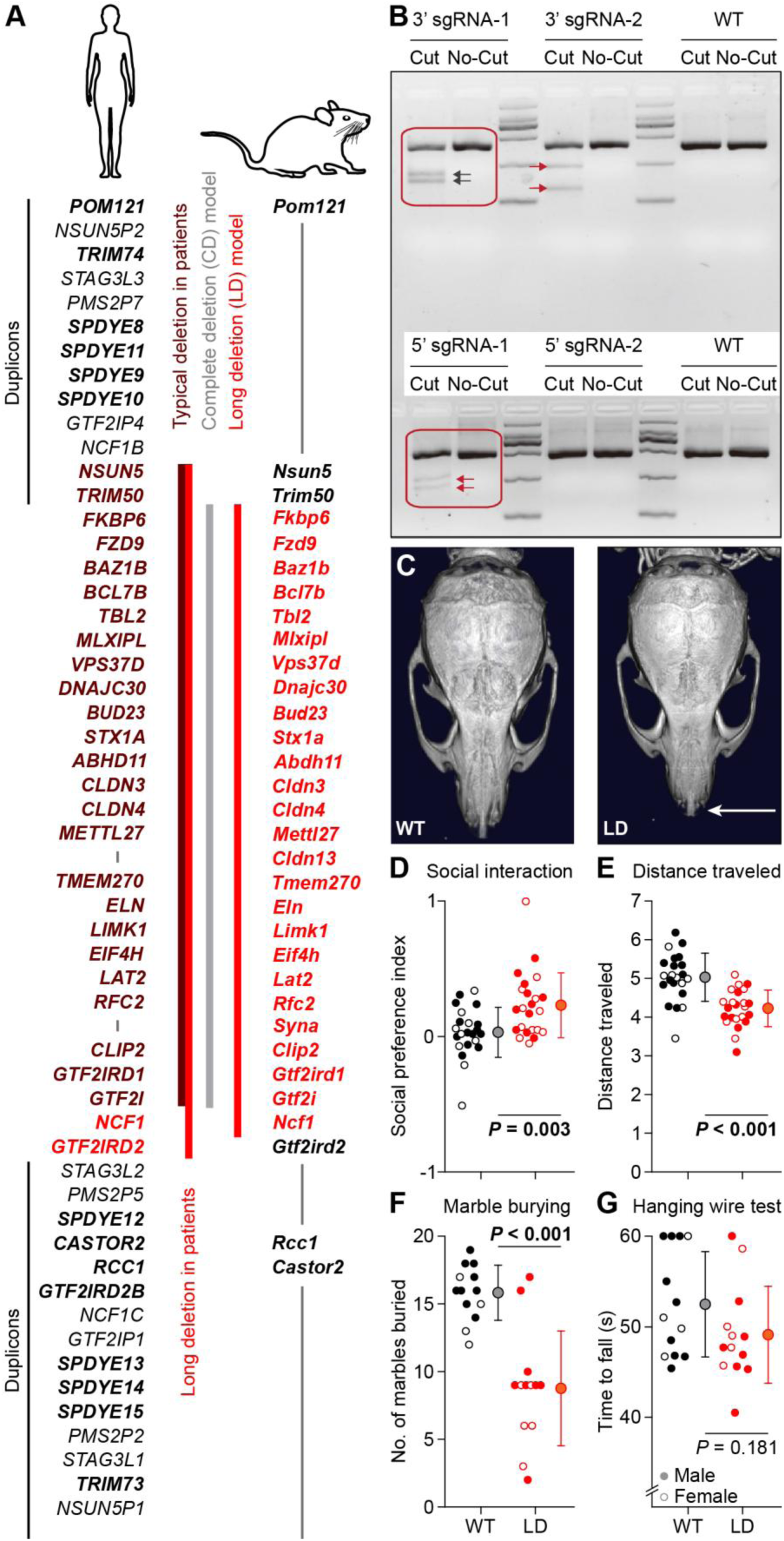
Creation and validation of the LD mouse model for Williams syndrome. *(A)* Left: The Williams syndrome critical region (WSCR) on human chromosome 7 with the typical deletion with 25 and longer variant with 27 genes, located between two areas containing duplicated genes (bold) and pseudogenes. Right: the homologous region of mouse chromosome 5, showing that the long deletion of the LD mouse includes also *Ncf1*, in contrast to the deletion of the previous CD mouse. *(B)* The T7E1 mismatch detection assay was performed to evaluate the cutting efficiency. *(C)* LD mice have a more compact skull, evident from a shorter nasal bone (arrow). µCT scans: Fig. S3. (*D*) During the first phase of the three-chamber test, LD mice traveled less than WT littermates (*n* = 22 WT and 23 LD mice, *P* < 0.001, *t* = 4.857). (*E*) LD mice showed higher sociability in the three-chamber test (*n* = 22 WT and 23 LD mice, *P* < 0.001, *t* = 4.857). Fig. S4A-D. *(F)* LD mice buried less marbles (*n* = 13 mice/genotype, *P* <0.001, Mann-Whitney test). Fig. S4E. (*G*) LD mice had normal performance on the hanging wire test (*n* = 13 mice/genotype, *P* = 0.181, Mann-Whitney test). Population statistics depicted as averages ± sd.

Cardiovascular abnormalities are the main cause of morbidity and mortality in Williams syndrome, affecting approximately 80% of patients (Collins et al., 2010; Parrish et al., 2020; Bozlak et al., 2023). Among these, supravalvular aortic stenosis (SVAS) is the most common one (Collins et al., 2010; Parrish et al., 2020; Bozlak et al., 2023). Other vascular anomalies include peripheral pulmonary artery stenosis and to a lesser extent coarctation of the aorta, ventricular septal defect, patent ductus arteriosus, subaortic stenosis, while also hypertrophic cardiomyopathy can occur (Collins et al., 2010; Das et al., 2014; Bozlak et al., 2023).

Hemizygosity of the elastin (*ELN*) gene, which encodes tropoelastin, is thought to be the principal cause of these cardiovascular symptoms. Patients with mutations restricted to the *ELN* gene exhibit cardiovascular abnormalities similar to those in Williams syndrome but lack among others the syndrome’s characteristic facial features (Urbán et al., 2000; Terashita et al., 2019; Zhou et al., 2022). During development, vascular smooth muscle cells produce multiple isoforms of tropoelastin via alternative splicing. These isoforms are secreted into the extracellular matrix and extensively cross-linked to form a network (Belknap et al., 1996). This network not only provides arteries with elastic properties, but also contributes to efficient blood propulsion (Cocciolone et al., 2018).

The neurological and psychological symptoms of Williams syndrome have largely been attributed to hemizygosity of the transcription factors *GTF2I*, *GTF2IRD1* and *LIMK1* (Borralleras et al., 2015; Kozel et al., 2021; Vernerova and Solc, 2024) and structural proteins like *CLIP2* (Hoogenraad et al., 2002; Hoogenraad et al., 2004; van Hagen et al., 2007). The larger cerebral blood vessels of patients can appear tortuous and may exhibit stenoses (Williams et al., 1961; Wollack et al., 1996; Wint et al., 2014). However, a possible direct contribution of abnormalities of the vascularization of the nervous system to neuropsychological symptoms has received little attention. Investigating this has been challenging due to the predisposition for hypertension associated with cardiac hypertrophy in many patients (Kececioglu et al., 1993; Das et al., 2014) and to the fact that the commonly used animal models lack deletion of the *Ncf1* gene, which may influence hypertension risk (Del Campo et al., 2006; Kozel et al., 2014).

To study the relationship between the WSCR deletion and vascular abnormalities in both the heart and brain, we generated a new mouse model carrying a long deletion (LD) that includes *Ncf1* (Fig. 1A). In contrast to other comprehensive mouse models of Williams syndrome (Segura-Puimedon et al., 2014), this LD model lacks a genetic driver of hypertension and more closely mimics the 1.85 Mb deletion found in a subgroup of patients (Del Campo et al., 2006; Kozel et al., 2014). Our model faithfully recapitulates the symptom spectrum observed in patients, but without the confounding effects of hypertension-induced cardiac hypertrophy. As such, this LD model offers a valuable tool to investigate the molecular, anatomical, and functional consequences of the WSCR microdeletion.

Notably, our LD model shows structural abnormalities not only in large vessels but also in small arterioles within the brain. Furthermore, the LD mouse model also reveals novel structural abnormalities in the inferior colliculus and cerebellum, which are consistent with auditory and motor phenotypes reminiscent of patient symptoms.

## Results

### LD mice reproduce key features of Williams syndrome

To generate a mouse model with a long deletion (LD) affecting the genes between *Trim50* and *Gtf2ird2*, we employed CRISPR/Cas9-mediated excision (Fig. 1A-B and Fig. S1). After modifying the site-specific locations of the selected genes using CRISPR/Cas9, two cuts were introduced on either side of the region of interest on chromosome 5, the mouse homolog of human chromosome 7q11.23 (Bayés et al., 2003; Kozel et al., 2021). A single-stranded oligodeoxyribonucleotide was inserted to ligate the two free ends of the excised region. In total, 48 mouse lines were generated on a C57BL/6J background and tested for DNA excision efficiency. Based on these tests, one line was selected and bred for further analyses. Heterozygous mice from this line were viable and were designated LD mice.

Our new LD mice showed a mildly reduced body weight (Fig. S2) and reproduced key anatomical and behavioral features of Williams syndrome, as well as of previous mouse models (Table S1). First, micro-computed tomography (µCT) revealed cranial abnormalities in LD mice, particularly a shorter nasal bone and a flattened parietal bone (Fig. 1C and Fig. S3), as found in *Gtf2ird1*-null and CD mouse models (Tassabehji et al., 2005; Segura-Puimedon et al., 2014) as well as in patients (Williams et al., 1961; Liu et al., 2021). Second, LD mice showed altered performance in tests for social and obsessive behavior (Fig. 1D-F), as typically found in patients and mouse models of Williams syndrome (Doyle et al., 2004; Kopp et al., 2019; Ortiz-Romero et al., 2021; Vernerova and Solc, 2024). LD mice had an increased tendency to visit the chamber with another mouse during the three-chamber test (*P* = 0.003, Mann-Whitney test; Fig. 1D and Fig. S4A-B). This increased sociability was not associated with other deviant behavior during the three-chamber test, except for a decreased distance traveled when exploring the setup during the first phase, i.e., before the second mouse was present (*P* <0.001, Mann-Whitney test, Fig. 1E and Fig. S4C-D). These results are consistent with increased sociability observed in other mouse models for Williams syndrome (Li et al., 2009; Segura-Puimedon et al., 2014; Kopp et al., 2019; Ortiz-Romero et al., 2021) and with those obtained with the use of another method for measuring social behavior, the modified elevated gap interaction test (De Zeeuw et al., 2024). LD mice buried fewer marbles (WT: 16 [2]; LD: 9 [3]; medians [interquartile range (IQR)]; *P* <0.001, Mann-Whitney test, *n* = 13 mice/genotype; Fig. 1F and Fig. S4E), suggesting increased behavioral flexibility. The hanging wire test, to assess gross motor deficits, did not reveal obvious differences between WT and LD mice (Fig. 1G). Overall, we conclude that the LD mouse model recapitulates key facial and neuropsychological symptoms observed in patients with Williams syndrome.

### Aortic abnormalities in LD mice

Given that many patients with Williams syndrome present with aortic abnormalities (Collins et al., 2010; Parrish et al., 2020; Bozlak et al., 2023), we next examined the 3D morphology of the aortas using µCT. LD mice have longer and more tortuous ascending aortas with a smaller diameter compared to WT littermates (Fig. 2A-B and Fig. S5A). Ultrasound imaging revealed reduced aortic distensibility in LD mice, indicating increased aortic stiffness (Fig. 2C-D and Fig. S5B). Histological analysis of the ascending aorta showed disrupted and disorganized elastic lamellae, along with an increased number of elastic lamellae (Fig. 2E-F). These abnormalities were associated with an elevated number of cells in the aortic wall (Fig. S5C-D). Since vascular malfunction is frequently linked to extracellular matrix remodeling, we assessed matrix metalloproteinases (MMPs) activity (Wågsäter et al., 2011). Using MMPSense 680 as a fluorescent probe, we did not find increased MMP activity of aortas in LD mice – the fluorescent signal was actually lower in LD mice (Fig. S5E-F). Nevertheless, a lack of elasticity of the aorta is also suggested by analysis of cardiac function with echocardiography, revealing an increased ejection fraction and fractional shortening in LD mice, indicative of enhanced cardiac contractility (Fig. 2G-H and Fig. S6). These findings suggest a compensatory cardiac response to the altered aortic structure.

**Figure 2.**
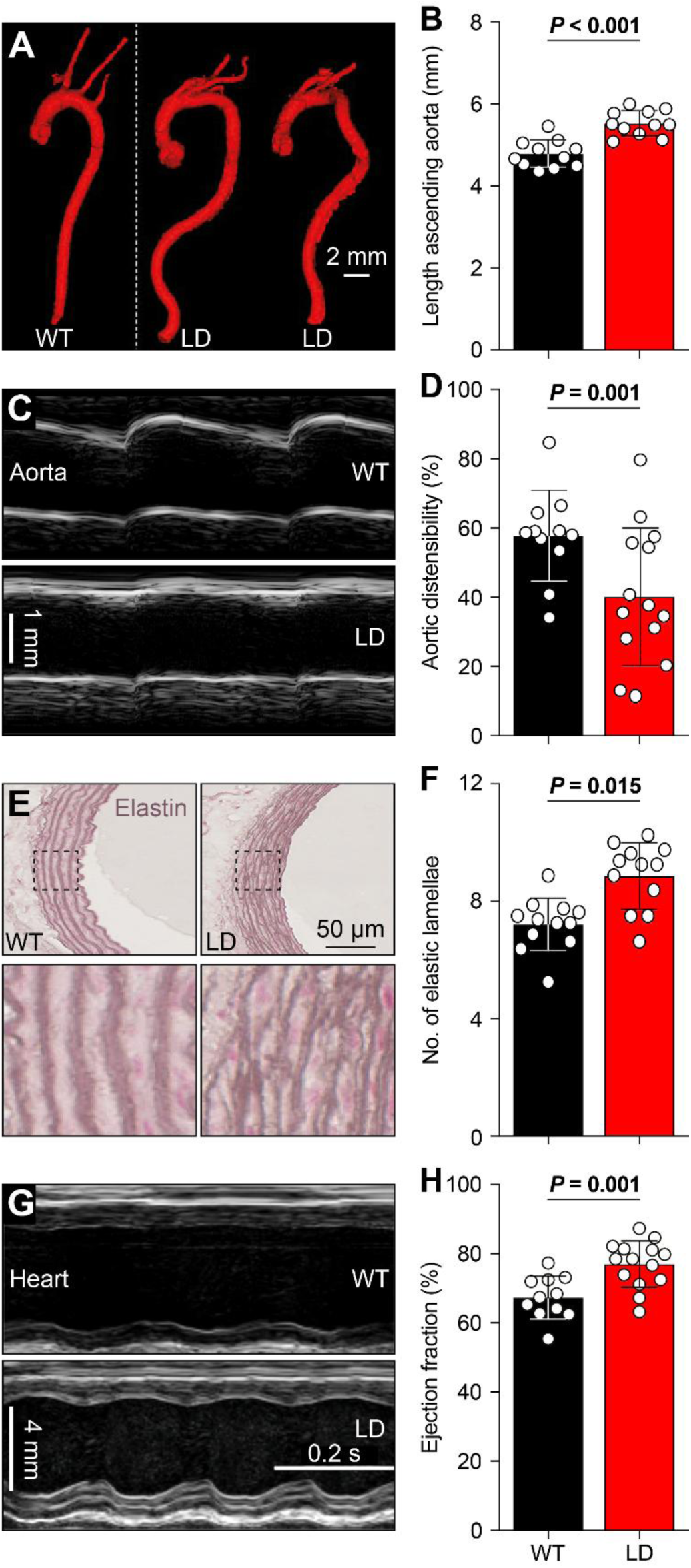
Tortuous and stiff aortae are associated with increased cardiac function in LD mice. (*A*) LD mice have a more tortuous aorta than WT littermates. *(B)* The ascending part of the aorta of LD mice is elongated (*n* = 11 mice/genotype, *P* < 0.001, *t* = 5.400, *t* test). *(C)* Representative M-mode images showing less expansion of the aorta during systole in LD mice (see also Fig. S5B). *(D)* This distensibility was indeed less than in WT mice, indicating reduced elasticity of the aortic wall (*n* = 11 WT and 14 LD mice, *P* = 0.014, *t* = 2.663, *t* test). *(E)* Resorcin-fuchsin stainings of cross-sections of the aorta showing more, but disorganized elastic lamellae in LD mice (*F*) Quantification of the elastic lamellae (*n* = 12 WT and 12 LD mice, *P* = 0.015, Mann-Whitney test). (*G*) Representative M-mode images from WT and LD mouse hearts demonstrating increased cardiac function in LD mice. (*H*) Increased ejection fraction in LD mice (*n* = 11 WT and 14 LD mice, *P* = 0.001, Mann-Whitney test). Population statistics depicted as averages ± sd.

### Normal cardiac morphology despite altered coronary arteries

In *Eln*^+/-^ mice, aortic wall changes are linked to hypertension and cardiac hypertrophy (Hawes et al., 2020). In contrast, LD mice do not display abnormal heart weight upon autopsy, nor was there an increase in cardiomyocyte size as indicated by FITC-labelled wheat-germ-agglutinin (WGA) staining. Accordingly, hearts of LD mice did not show any myocyte disarray or significant differences of interstitial fibrosis (Fig. S7A-D). Thus, despite altered cardiac function, no structural remodeling was observed in the hearts of LD mice.

We next examined the coronary arteries using Microfil casting combined with µCT. Similar to the aorta, coronary arteries in LD mice were longer and more tortuous, resulting in increased coronary volume (*P* = 0.039, *t* test, Fig. 3A-B), as also seen in patients (Pober et al., 2008). In contrast to in the aorta, we found increased MMP activity in the coronary arteries of LD mice (Fig. 3C-D). These findings suggest coronary remodeling in the absence of overt cardiac hypertrophy.

**Figure 3.**
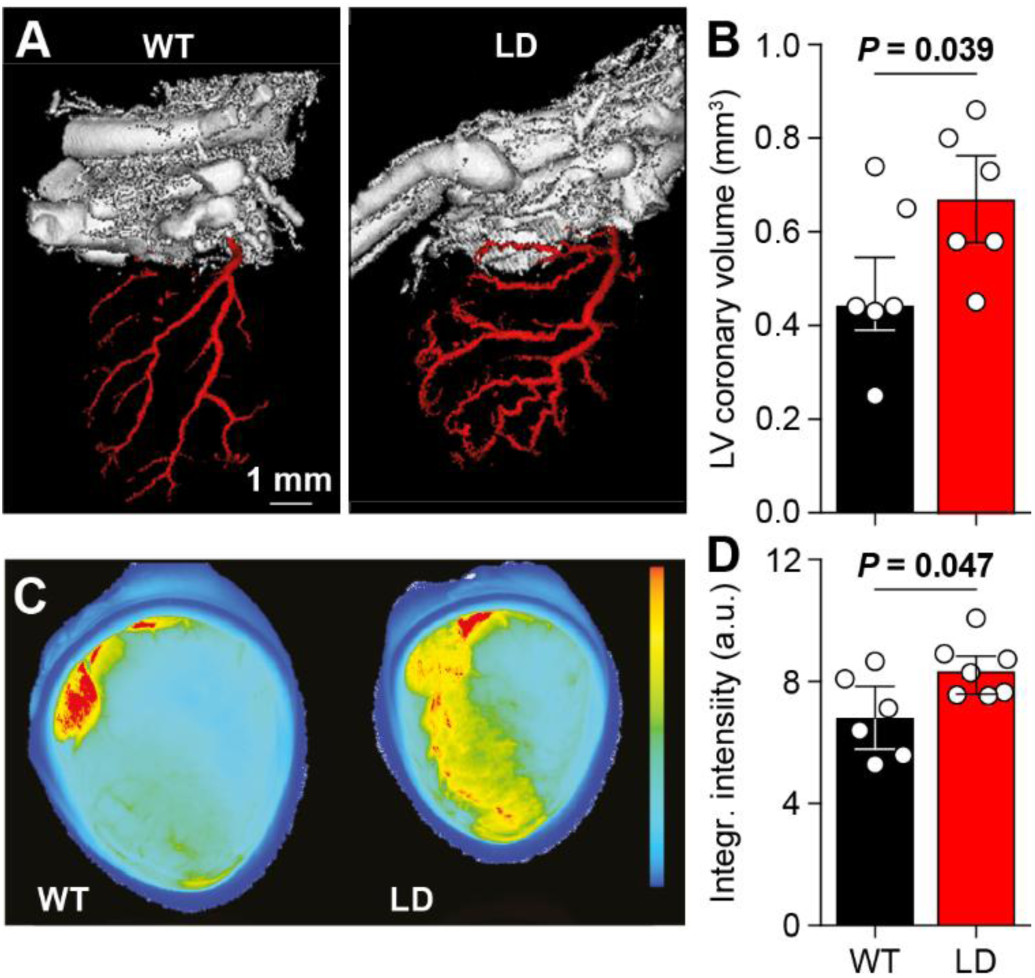
Coronary arteries are tortuous and remain plastic in LD mice. (*A*) 3D rendering of the coronary arteries using Microfil blood vessel casting in combination with µCT imaging. (*B*) Total left ventricle coronary volume (*n* = 6 mice/genotype, *P* = 0.039, *t* = 2.345, *t* test). *(C)* Representative scans showing elevated levels of matrix metalloprotein (MMP) activity in hearts of LD mice, indicating increased matrix remodeling of the cardiac coronary arteries. (*D*) Fluorescence due to MMP activity (*n* = 6 WT and 7 LD mice, *P* = 0.047, *t* = 2.348, *t* test). Population statistics depicted as medians and inter-quartile range.

### Alterations in cerebral vasculature

Unlike the aortic and coronary arteries, cerebral vascular abnormalities in animal models for Williams syndrome have been relatively underexplored, although case reports indicate tortuosity and stenosis of major cerebral vessels in patients (Williams et al., 1961; Soper et al., 1995; Wint et al., 2014). To assess cerebral vascular changes in LD mice, we first used post mortem µCT to evaluate the circle of Willis and other major vessels, which revealed increased tortuosity (Fig. 4A and Fig. S8).

**Figure 4.**
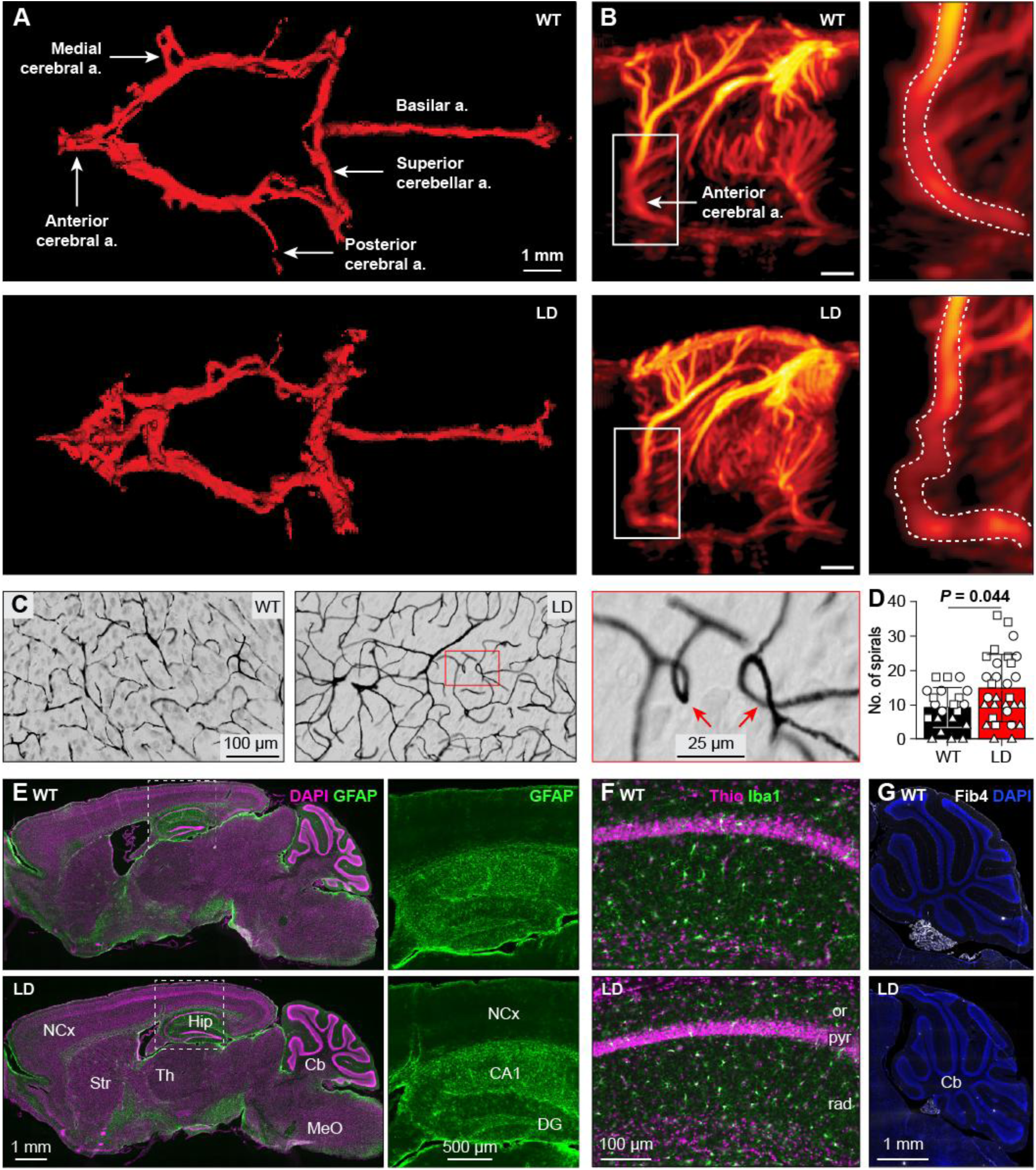
Abnormal vasculature in the brain. *(A)* Representative µCT images of the circle of Willis and related blood vessels. *(B)* In vivo ultrafast Doppler imaging of the cerebral vasculature in two example mice. See also Fig. S8-10. *(C)* Brain microvasculature visualized after perfusion with Indian ink. LD mice had more spirals, as exemplified by zoom image. (*D*) Quantification of the number of spirals after counting them in standardized areas of the hippocampus (triangles), cerebellar cortex (circles) and medulla (squares) (*n* = 7 WT and 10 LD mice, *P* = 0.044, *F* = 4.824, repeated measures ANOVA). Average ± sd. (*E*). Unaltered GFAP (glial fibrillary acidic protein) staining in the brains of LD mice. Cb = cerebellum, Hip = hippocampus, NCx = neocortex, MeO = medullar oblongata, Str = striatum, Th = thalamus. (*F*) Normal distribution, number and size of microglia in the brains of LD mice as visualized with Iba1 immunofluorescence. or = stratum oriens, pyr = pyramidal layer, rad = stratum radiatum. (*G*) No upregulation of Fibulin-4 (Fib4) in the brains of LD mice.

To assess smaller vessels, we used in vivo ultrafast Doppler imaging, which confirmed increased cerebral vessel tortuosity in LD mice (Fig. 4B and Fig. S9). For the analysis of the microvasculature, we performed ink perfusion followed by histological processing of 50 µm-thick brain sections. LD mice exhibited more irregular vascular structures, including stubs, right-angled bends and spirals (Fig. 4C and Fig. S10). Quantification of spirals in the cerebellum, hippocampus and brainstem showed a significant increase in LD mice (*P* = 0.044, *F* = 4.824, repeated measures ANOVA, Fig. 4D). Combined, these findings indicate that both large and small cerebral vessels are morphologically altered in LD mice.

As abnormalities in the cerebral microvasculature can be associated with inflammation (Gama Sosa et al., 2023), we examined glial responses. GFAP immunoreactivity showed no changes in astrocytes (Fig. 4E), and IBA1 staining indicated no alterations in microglia number or morphology (Fig. 4F). Similarly, Fibulin-4 immunoreactivity, often associated with vascular pathology (Stefens et al., 2024), was not upregulated in neural tissue of LD mice (Fig. 4G). We, therefore, conclude that the changes in microvasculature in the brains of LD mice probably reflect a primary genetic defect, rather than a secondary compensation or maladaptation to inflammatory processes.

### Alterations in brain anatomy and function

While examining brain slices, we noticed alterations in the shape of the brain, in particular of the morphology of the inferior colliculus and cerebellar vermis. Indeed, the size of the inferior colliculus was reduced in LD mice, whereas that of lobule IV/V of the cerebellar vermis was significantly enlarged (Fig. 5A-C and Fig. S11A). Given that these two structures are adjacent to each other, this configuration raises the possibility that the cerebellar enlargement occurred at the expense of the inferior colliculus.

**Figure 5.**
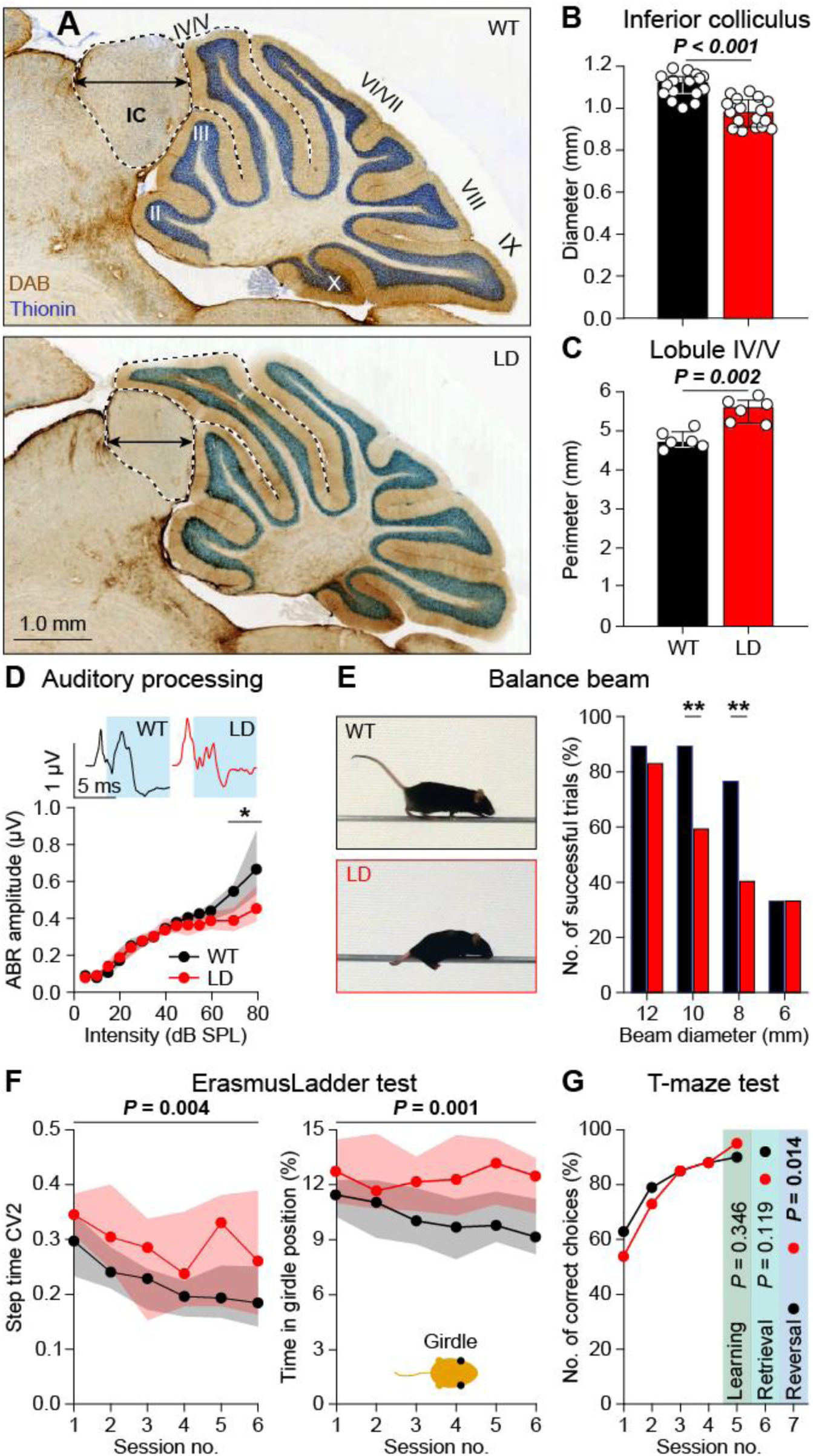
Sensory, motor and cognitive symptoms in LD mice. (*A*) LD mice had a smaller inferior colliculus (IC) and an expanded cerebellar lobule IV/V. (*B*) Decreased diameter (arrow in *A*) of the inferior colliculus in LD mice (*n* = 16 WT and 17 LD mice, *P* < 0.001, *t* = 6.002, *t* test). (*C*) The perimeter (dotted black line in *A*) of cerebellar lobule IV/V was larger in LD mice (*n* = 6 mice/genotype, *P* < 0.002, Mann-Whitney test). (*D*) Representative auditory brainstem response (ABR) in response to an 80 dB SPL, 12 kHz tone. LD mice showed abnormal ABR to tones >50 dB SPL (*n* = 14 WT and 11 LD mice, *P* = 0.009, repeated measures ANOVA). (*E*) With intermediate beam diameters, LD mice were less able to walk across the balance beam, and tended instead to crawl (*n* = 13 WT and 14 LD mice, *P* = 0.522 (12 mm), *P* = 0.002 (10 mm), *P* = 0.002 (8 mm), *P* = 1.000 (6 mm), Fisher’s exact test, ** *P* < 0.01 after Bonferroni correction for multiple comparisons). (*F*) On the ErasmusLadder, LD mice had a more variable stepping pattern (step time CV2: *n* = 21 mice/genotype, *P* = 0.004, *F* = 9.577, repeated measures ANOVA). When stepping on the rungs, the LD mice were more often found in the girdle position (thus, with both front limbs touching the rungs; *P* = 0.001, *F* = 11.679, repeated measures ANOVA). (*G*) Both WT and LD mice rapidly learned to choose the arm with a shelter on a T maze (*n* = 18 WT and 15 LD mice, session 5 (end of learning): *P* = 0.346, session 6 (retrieval after 1 week): *P* = 0.119, session 7 (target arm reversal): *P* = 0.014 (significant after Bonferroni correction for multiple comparisons), Fisher’s exact tests). Population statistics depicted as medians with inter-quartile range.

As the inferior colliculus is crucial for auditory processing, we wondered whether hearing could be affected in LD mice. Hence, we examined auditory brainstem responses to pure tones. In line with data from other mouse models for Williams syndrome (Canales et al., 2015; Davenport et al., 2022), we found that the hearing thresholds and early responses (peak I) were unaltered (Fig. S12A-C, I). However, later responses (peaks IV-VI) were significantly affected at high sound intensities (Fig. 5D and Fig. S12). These observations suggest that, even though auditory nerve activity was relatively normal, central processing of loud sounds at the level of the inferior colliculus and higher auditory centers could be altered in LD mice. These data are compatible with the symptoms of patients with Williams syndrome, who can suffer from hyperacusis and progressive hearing loss (Klein et al., 1990; Gothelf et al., 2006).

The enlargement of cerebellar lobule IV/V in LD mice was rather specific (Fig. 5C), as none of the other lobules in the vermis, hemispheres or vestibulocerebellum showed aberrant volumes (Fig. S11B). Given that lesion and recording studies indicate that lobule IV/V is involved in motor coordination (Hoogland et al., 2015; Chao et al., 2021), we first tested the mice on the balance beam. LD mice were less able to walk fluently from one side to the other under challenging conditions, i.e., if the beam diameter was relatively small (Fig. 5E). This phenotype was reflected in the number of successful trials rather than in the average crossing time, because when unable to walk, the LD mice often crawled across the beam with a speed that was comparable to walking (Fig. S13).

To characterize the gaiting pattern, we next performed locomotion tests on the ErasmusLadder, which consists of a horizontal ladder with alternating higher and lower rungs between two shelter boxes (Fig. S14A). As on the balance beam, travel times on the ErasmusLadder were unaltered in LD mice (Fig. S14B-D). However, LD mice more often made small steps, which was statistically significant also when taking their reduced body weight into account (*P* = 0.012, *F* = 6.913, repeated measures ANOVA, Fig. S14E-G).

The number of lower rung touches, which is typically increased in mouse models for cerebellar ataxia (Vinueza Veloz et al., 2015; Jaarsma et al., 2024), was unaltered in LD mice (Fig. S14H). However, we noticed a more subtle sign of ataxic gait, as the variation (CV2) in step times was increased (*P* < 0.004, *F* = 9.577, repeated measures ANOVA, Fig. 5F). The increased variation in gaiting pattern motivated us to further analyze the inter-limb coordination, which is also typically affected by cerebellar deficits (Jaarsma et al., 2024), to note that LD mice more often use the girdle position, thus touching the rungs with their two front limbs (*P* = 0.001, *F* = 11.679, repeated measures ANOVA, Fig. 5F and Fig. S14I). Indeed, the interlimb coordination of LD mice was affected in that we observed a noticeably higher level of variability for the movements across all body axes using an unsupervised machine learning analysis (Fig. S15). Thus, LD mice do not suffer from gross deficits in motricity, but they indeed do have an impairment in motor coordination of limb movements (Fig. S14J-K).

Another pattern we observed on the ErasmusLadder was a slightly different distribution over time in the percentage of early escapes from the starting box, with the LD mutants showing an increased number of early escapes in the later sessions (Fig. S16). Akin the outcome of the marble burying test (Fig. 1E), this phenotype on the Erasmus Ladder may reflect different levels of motivation or fear as well as related emotional memory formation thereof (Vinueza Veloz et al., 2015). We, therefore, tested LD mice on a T-maze to find out whether they were flexible in learning and relearning targets in space. Whereas their memory retention after 1 week was similar to that in WTs, LD mice were faster in switching their behavior when we changed the target arm on the last day (*P* = 0.014, Fisher’s exact test, Fig. 5G and Fig. S17). Thus, the flexibility of LD mice to adjust their behavior was apparently not affected by high levels of fear, in line with the lower number of marbles buried as well as with the high number of early escapes in the later sessions on the ErasmusLadder.

## Discussion

Williams syndrome is a complex heterozygous genetic condition. Although atypical forms exist, most patients have a deletion affecting 25 genes, while a longer deletion including 27 genes occurs in a sizeable subpopulation (Peoples et al., 2000; Bayés et al., 2003; Fusco et al., 2014; Lugo et al., 2020; Kozel et al., 2021). Both most prevalent forms of Williams syndrome include *ELN* deletion, but *NCF1* is only affected in the longer deletion. This is of interest as there is a gene dosage effect of *NCF1* on the manifestation of cardiovascular symptoms in patients with Williams syndrome (Kozel et al., 2014). To study the impact of *Ncf1* inclusion in the deletion causing Williams syndrome, we created a novel mouse model with a long deletion (LD), contrasting the previously created CD mouse model (Segura-Puimedon et al., 2014) (Fig. 1A). The LD model replicates major craniofacial and neurological symptoms of patients, including hypersocial behavior, altered auditory sensitivity, and a mild motor deficit, as well as the typical vascular deficits, including SVAS, aortic stiffness and tortuosity. Despite altered cardiac function, cardiac pathology with hypertrophy was absent in the LD model, in contrast to the CD model (Segura-Puimedon et al., 2014), hinting at gene interactions. Despite the absence of cardiac hypertrophy, our LD model for Williams syndrome did show abnormal vascularization of not only the heart, but also of the brain. This finding raises the possibility that the neuropsychological symptoms in Williams syndrome are not only caused by haploinsufficiency of genes like *GTF2I*, *GTF2IRD1*, *LIMK1* and *CLIP2* (Borralleras et al., 2015; Kozel et al., 2021; Vernerova and Solc, 2024), but also by that of the WSCR gene(s) implicated in structure and function of the vasculature.

### Roles of ELN and NCF1

It is generally assumed that haploinsufficiency of *ELN* is the main culprit for the vascular phenotype in Williams syndrome (Kozel et al., 2021; Ganjibakhsh et al., 2025). Indeed, patients with loss-of-function mutations in *ELN* develop elastin arteriopathy, but not the craniofacial or neurological symptoms of Williams syndrome (Li et al., 1997; Tassabehji et al., 1997; Min et al., 2020; Stephens et al., 2024; Ganjibakhsh et al., 2025). Just as in Williams syndrome, patients carrying loss-of-function mutations confined to *ELN* often present with SVAS, thickened aortic walls and reduced aortic lumina (Li et al., 1997; Tassabehji et al., 1997; Phomakay et al., 2015; Min et al., 2020; Stephens et al., 2024; Ganjibakhsh et al., 2025). This has been replicated in mice that are haploinsufficient for *Eln* (Faury et al., 2003; Hirano et al., 2007; Knutsen et al., 2018; Hawes et al., 2020). Remarkably, in patients with non-syndromic SVAS, surgical intervention is often required at an earlier age, and valvar problems are more abundant than in Williams syndrome (Min et al., 2020). Thus, despite the overlap in vascular deficits between isolated *ELN* loss-of-function mutations and Williams syndrome, the symptoms are not identical, indicating that other genes in the WSCR may modulate the phenotype.

In this respect, the *NCF1* gene at the telomeric end of the WSCR may be relevant. Inclusion of *NCF1* in the deletion, which occurs in a subgroup of patients with Williams syndrome, reduces the prevalence of cardiac hypertrophy with hypertension (Del Campo et al., 2006; Kozel et al., 2014; Phomakay et al., 2015). Reduced expression of *NCF1* can limit the impact of *ELN* haploinsufficiency on hypertension by ameliorating the increase of reactive oxygen species in the vascular walls through regulation of NOX1 and NOX2 NADPH oxidase complexes (Del Campo et al., 2006; Kozel et al., 2014; Troia et al., 2021; Abdalla et al., 2023). The same mechanism may explain why our LD mouse model replicated many but not all of the phenotypes observed in the CD model; both models showed more chaotic elastic lamellae, increased smooth muscle cells in the vessel walls, and a reduced aortic lumen (Segura-Puimedon et al., 2014; Jiménez-Altayó et al., 2020), but the hearts of LD mice did not show the hypertrophic pathology observed in CD mice (Segura-Puimedon et al., 2014; Ortiz-Romero et al., 2018). As the only difference between the CD and LD mouse models is the inclusion of *Ncf1*, WSCR genes may indeed interact; *NCF1* deletion may partially compensate for the haploinsufficiency of *ELN* and thereby influence the pathological processes leading to cardiac problems in Williams syndrome. The variation in clinical manifestations of Williams syndrome, with different levels of abnormalities in ejection fraction and cardiac hypertrophy (Deitch et al., 2023), further underscores the need to understand the relationship between genetic factors and disease severity.

### Vascular abnormalities in heart and brain

In line with vascular deformations seen in patients with Williams syndrome (Williams et al., 1961; Soper et al., 1995; Wint et al., 2014), abnormalities of the vessels were more common in LD than in WT mice, affecting all types of vessels and ranging from the aorta to capillaries, and from heart to brain. The larger vessels in heart and brain appeared similarly affected, showing severe irregularities, including changes in diameter, length, and shape. However, with respect to smaller vessels there were slight differences between heart and brain in that deformations of smaller vessels in the heart, but not in the brain, could be associated with abnormalities in the surrounding tissue in LD mice. Indeed, the increased tortuosity of the coronary arteries in LD mice occurred alongside with increased levels of MMP activity, indicative of remodeling due extracellular matrix degradation (Hutchinson et al., 2010; Wågsäter et al., 2011; Brkic et al., 2015; Bräuninger et al., 2023). In contrast, the increased vascular tortuosity of smaller brain vessels and the high density of spirally shaped capillaries were not associated with signs of degradation or degeneration in the extra-neuronal space. For example, the structure and function of astrocytes and microglia appeared unaffected in LD mice.

Despite the potentially differential responses of heart and brain to similar alterations in vascularization, it should be noted that increased tortuosity is associated with ischemia not only in the heart but also in the brain (Han, 2012; Ciuricã et al., 2019). Hence, cerebral arteriopathy may be another mechanism contributing to the neuropsychological symptoms of Williams syndrome, alongside the putative impact of transcription factors such as *GTF2I* and *LIMK1*, and structural proteins like CYLN2 (Tassabehji et al., 1999; Hoogenraad et al., 2002; Hoogenraad et al., 2004; van Hagen et al., 2007; Borralleras et al., 2015; Kozel et al., 2021).

### Novel structure-function correlations in LD mice

Morphological and functional analyses of LD mice revealed two associations not previously described in animal models of Williams syndrome: changes in the volume of the inferior colliculus and cerebellar vermis, which may relate to altered auditory processing and motor coordination, respectively. These novel relations may be relevant for patients, who often suffer from both hyperacusis and mild motor deficits (Klein et al., 1990; Don et al., 1999; Gothelf et al., 2006; Meyer-Lindenberg et al., 2006).

Hyperacusis is very common in patients with Williams syndrome and many patients show increased affinity for music, liking or disliking specific sounds and melodies (Klein et al., 1990; Don et al., 1999; Gothelf et al., 2006; Meyer-Lindenberg et al., 2006). Children with atypical deletions sparing *GTF2I* are less prone to develop hyperacusis (Tassabehji et al., 1999; van Hagen et al., 2007). In LD mice, we observed normal responses of the auditory nerve and cochlear nucleus, but not at higher auditory centers. Altered auditory processing was accompanied by hypoplasia of the inferior colliculus. The functional differences were most apparent at higher sound intensities (>60 dB SPL). This differential processing of loud sounds may explain why hyperacusis was not noted in previous mouse models for Williams syndrome (Canales et al., 2015; Davenport et al., 2022), despite improved inhibition of the acoustic startle response and increased auditory acuity (Li et al., 2009; Canales et al., 2015; Davenport et al., 2022). The observation that peak IV is altered in LD mice points to a neural basis for hyperacusis at the level of secondary auditory centers, including the inferior colliculus (Henry, 1979; Land et al., 2016)

Likewise, mild motor deficits are common in patients with Williams syndrome (Meyer-Lindenberg et al., 2006; Kozel et al., 2021). In LD mice, both the balance beam and ErasmusLadder tests showed impaired inter-limb coordination during gait, and shorter steps. These symptoms are in line with relatively subtle cerebellar deficits (Vinueza Veloz et al., 2015; Jaarsma et al., 2024). Strikingly, we observed hyperplasia of vermal lobule IV/V in LD mice, reminiscent of findings in patients (Reiss et al., 2000; Schmitt et al., 2001). Given that vermis lobule V is involved in locomotor coordination (Hoogland et al., 2015; Chao et al., 2021), it is conceivable that the mild motor deficits in LD mice result at least partly from malformation of their cerebellar vermis. Thus, even though further mechanistic studies have to be done on the inferior colliculus and vermis of the cerebellum in LD mice to demonstrate a causal role in the abnormalities in hearing and locomotion, these novel phenotypes are remarkably similar to the symptoms observed in patients, both at the structural and functional level.

### Flexible and social behavior in LD mice

Patients with Williams syndrome often show a mosaic IQ pattern, with relatively normal language and social skills (Don et al., 1999; Meyer-Lindenberg et al., 2006). Indeed, given their low IQ, WS patients appear rather flexible and hypersocial (Doyle et al., 2004; Vernerova and Solc, 2024). LD mice confirm this behavioral pattern in that they showed a relatively flexible behavior in the marble burying test, similar to what has been found for CD mice (Kopp et al., 2019; Ortiz-Romero et al., 2021). The increased level of flexibility was underscored in the Erasmus Ladder test, where they often left the starting box early during the later sessions, in line with low levels of fear (Vinueza Veloz et al., 2015). Moreover, during the T-maze task LD mice were also more flexible in relearning new targets in space than the WT littermates.

The high level of flexibility of LD mice was also apparent in social settings. LD mice showed a stronger bias toward the social chamber in the three-chamber test, which is in line with findings done in other WS mouse models (Li et al., 2009; Segura-Puimedon et al., 2014; Kopp et al., 2019; Ortiz-Romero et al., 2021). Moreover, they also appear hypersocial when directly interacting with juvenile mice in the modified elevated gap interaction test (De Zeeuw et al., 2024). Overall, we conclude that relatively high levels of flexibility and an eagerness for social interactions form hallmarks of the Williams syndrome spectrum, both in the patients and the LD mouse model.

## Materials and Methods

### Ethics Statements

Animals were generated and the experiments were performed at the Chinese Academy of Science (CAS) according to institutional guidelines as overseen by Animal Welfare Board of the CAS, following Chinese legislation, as well as at the Erasmus MC, following Dutch and EU legislation. Prior to the start of the experiments, project licenses for the animal experiments performed for this study at the Erasmus MC was obtained from the Dutch national authority and filed under no. AVD101002015273, AVD1010020197846 and AVD10100202216286

### Generation of the LD mouse model

To generate the LD mouse model, CRISPR/Cas9 technology was used (Leonova and Gainetdinov, 2020) to delete the genes between *Trim50* and *Gtf2ird2* on chromosome 5. The genes within this region correspond to the analogous region on chromosome 7q11.23 in humans (DeSilva et al., 2002), whose heterozygous deletion leads to the development of Williams syndrome. CRISPR/Cas9 technology modifies the site-specific locations of the selected genes, leading to deletions of this region. Two cuts were performed on each side in our chromosome region of interest. The knockout fragment equals 1.1 Mb. After cutting, filaments of single-stranded oligodeoxyribonucleotide (ssODN) (100nt) were inserted on each side of the region of the deleted gene to attach the two free extremities of each cut region. The sequences of the single guide RNA (sgRNA) are: 5’ sgRNA-1 is agggccacacccatggctgc; for 5’ sgRNA-2 is ctgagtgcacatatagtgct; 3’ sgRNA-1 is tcccaaggtcatccagaatc; 3’ sgRNA-2 is ggtcacacaccacgagactg. After that, a T7 endonuclease 1 (T7E1) mismatch detection assay was performed to evaluate the cut efficiency or the activity of site-specific nucleases (Sentmanat et al., 2018), and this revealed a high cut efficiency (Fig. 1B). In total, 48 mice were generated and tested for their cutting DNA efficiency. Finally, Sanger sequencing of polymerase chain reaction product (PCR) was performed to confirm the real cutting. The mice had a C57BL/6J background.

Animals were group housed in a general mouse facility. Environment was controlled with a temperature of 20–22°C and 12 h light:12 h dark cycles. Food and water were offered ad libitum. Experiments were performed on groups of mixed sex. Experimenters were blind to the genotype of the mice during the experiments and initial analyses.

### Auditory brainstem responses

LD mice and WT littermates (age: 87-92 days) were anesthetized using ketamine/xylazine (80 mg/kg, 8 mg/kg) and placed in front of an electrostatic speaker (ES1, Tucker-Davis Technologies, Alachua, FL) at 5-6 cm distance in a sound-attenuating chamber. Body temperature was monitored via a rectal probe, and maintained between 36.0-37.5 °C. Auditory brainstem responses were measured using a subcutaneous four-electrode configuration with the reference electrode at the vertex, two lead electrodes ventral of the ears and a ground electrode at the level of the caudal vertebrae. Tone pips of 4, 8, 12, 16 and 32 kHz and 1 ms duration were presented ranging from 5 to 80 dB sound pressure level (SPL, re 20 μPa). RZ6 multi-I/O processor (Tucker Davis Technologies) generated the tones. Recordings were pre-amplified (Medusa4Z, Tucker-Davis Technologies) and digitized by the RZ6. Hardware was controlled by BioSigRZ (Tucker-Davis Technologies). Recordings were bandpass filtered (10-5000 Hz), a minimum of 800 recordings were averaged, and then the waveforms of the two recording electrodes were averaged. Thresholds were defined as the lowest intensity at which a reproducible waveform could be identified by visual inspection. Amplitudes were measured within a time window set by the experimenter. In this window the difference between the maximal value and the minimal value preceding the maximal value was taken as the amplitude. After the ABR recordings, mice were transcardially perfused or euthanized.

### Behavioral testing

#### Hanging wire test

Mice were brought to a metal wire with a diameter of 2 mm that was suspended approximately 20 cm above a cage, so that they could grasp the wire with their front paws. Once a mouse grabbed the wire, the experimenter gently released the mouse and the latency to fall was recorded. The maximal trial duration was 60 s.

#### Balance beam

Mice were tested on the balance beam as described previously (Wahl et al., 2023). Briefly, the setup consisted of two platforms connected by a metal horizontal beam of 86 cm that was positioned 50 cm above an acrylic glass plate. Mice had to cross the beam from one platform to the other, after which, mice could enter their home cage. On the first training day, mice three times cross a beam with a diameter of 12 mm and three times on one with a diameter of 10 mm. On the second day, this is repeated, but with diameters of 10, 8 and 6 mm. On the third day, videos are made of three trials on each of the beams, starting with the thickest one.

The recorded crossings on the third day were analyzed on the middle 70 cm of the beams. The time to cross and the number of hindlimb slips were scored, and we noted whether the mice walked or crawled to the other side. Crossings during which the mice fell or jumped off the beam, or when they made a slip severe enough that they could not resume crossing without help from the experimenter, were considered as fails.

#### ErasmusLadder

Locomotor patterns were recorded on the ErasmusLadder (Noldus, Wageningen, the Netherlands), a horizontal ladder flanked by two plexiglass walls spaced 2 cm apart as described previously (Vinueza Veloz et al., 2015; Jaarsma et al., 2023). The ladder consists of two rows of 37 rungs spaced 15 mm apart and placed in an alternating high/low pattern. The height difference between high and low rungs is 9 mm. Each rung is connected to a pressure sensor recording rung touch.

During a trial, the mouse had to walk from a shelter box on one side of the ladder to another on the other end. Trial start was indicated by an LED in the shelter box followed three seconds later by a strong tail wind. Early escapes, thus before the LED was switched on, were discouraged by triggering a strong head wind. In between trials, there was a resting period. Mice were first habituated to the setup by letting them freely explore the ladder for 15 min during which no light or air cues were given. On the next day, training started with 42 trials on each day for five consecutive days. Only regular trials, thus trials without pauses or other disruptions, were analyzed as described previously (Jaarsma et al., 2023).

Hildebrand plots were made to visualize inter-limb coordination, as described previously (Jaarsma et al., 2024). Briefly, for each step we calculated the duty cycle as the percentage of the step time during with the right forelimb touched a rung and as the fraction of the step cycle between the placement of the right forelimb and a second limb. The number of steps were normalized per session, and, for each genotype and stance, we used unsupervised machine learning in combination with *k* means clustering to divide the histograms in two clusters. To this end, we used the pretrained InceptionV3 unsupervised machine learning algorithm from the Keras applications library in Python, whereby we removed the output layer of the pretrained model. The extracted image features were used as input for *k* means clustering (*k* = 2). To control for model sensitivity to the initial conditions, we ran each model ten times and compared the resulting clusters. The difference between the averages of groups 1 (“healthy”) and 2 (“affected”), were visualized as heat maps. Principal component analysis (PCA) of the ErasmusLadder data was performed on 23 different parameters, as described previously (Jaarsma et al., 2023).

#### Three-chamber test for social preference

Mice were placed in a box consisting of three chambers that were accessible by small doors in the two walls. The middle chamber was always empty. The test consists of five steps. In step 1, the mice were placed in the middle chamber for 5 minutes without access to the other chambers in order to habituate to the setup. In the second step (phase 1), all chambers were empty, and the baseline preference of the mice was tested for 10 minutes. For step 3 (phase 2), an empty wire mesh cup was placed in one of the outer chambers and a cup containing an age and sex matched mouse was placed in the other. The test mouse roamed freely for 10 minutes and preference for time spent with the object (empty cup) or conspecific was calculated (social preference). Step 4 consisted of placing the mouse in the chamber with the cup holding the other mouse for 5 minutes to make sure this mouse is familiar. For step 5 (phase 3), a second age and sex matched mouse was placed under the previously empty cup. During the next 10 minutes, preference for time spent with either the familiar or the novel mouse was calculated.

#### Marble burying test

Repetitive, obsessive-compulsive behavior was tested by placing the mice in a large cage with 20 marbles spaced evenly on top of the bedding for 30 minutes. Pictures were taken at 10, 20 and 30 minutes to assess the number of marbles buried. A marble was considered buried if less than 50% of the marble was visible.

#### T-maze test

Mice were place on one arm of a white T-maze under intense illumination. Both other arms had a hole (5 cm diameter), of which one was covered underneath the surface with a black plexiglass plate and the other was open, giving access to a shelter beneath the surface. The time to reach the entrance of any of the two arms was calculated from an overhead video (frame rate 30 Hz), as was the time to target area (defined as both ears within the opening) and target entrance (defined as all four limbs inside the shelter). After five daily sessions of 4 trials each, the mice were re-tested after 1 week. During the last session, on the day after the re-training, the mice started from the opposite arm, that was initially blocked by an opaque wall. As the target was still on the same side, the mice had to turn to make the opposite turn.

### Aortic and cardiovascular perfusion and µCT imaging

Prior to sacrifice, mice were given a sub-lethal i.p. dose of ketamine (100 mg/kg) plus xylazine (6 mg/kg). Vascular perfusions were carried out as described previously (Downey et al., 2012). The abdominal cavity was opened, the diaphragm was incised to expose the pleural cavity, and the ribs were cut away to allow access to the heart. A 25G butterfly needle attached to 0.75 in polyethylene tubing (BD Vacutainer) was inserted into the left ventricle and the right jugular vein severed to provide a drainage point. The circulatory system was flushed with 10 ml of heparinized saline, and this was immediately followed by perfusion with 10 ml of radio-opaque silicone rubber polymer (MicrofilH, MW-122 yellow, Flow-Tech) at a constant rate of 0.75 ml / min using a Harvard Apparatus PHD 2000 Infusion pump (Instech Laboratories Inc). The Microfil casting was allowed to polymerize for 20 min and then stored in formalin overnight at 4 °C. Hearts were excised and fixed in 10% neutral buffered formalin (NBF) for a minimum of 24 h prior to µCT scanning. Micro-CT imaging was performed using an ultra-fast µCT scanner (Quantum FX, PerkinElmer). The µCT acquisition parameters were 90 kVp, 160 µA with field of view 20 mm and 360° rotation in 1 step. The acquisition time was 4.5 min. Images in Hounsfield unit (HU) were reconstructed in cubic voxels of 40 µm, generating a matrix of 512 × 512 × 512 voxels. The bones and vascular network were visualized using the volume renderer available from MeVisLab (MeVisLab 2.2.1, MeVis Medical Solutions AG).

### Ultrafast Doppler imaging

Ultrafast Doppler imaging was used to characterize the brain vasculature. To this end, adult mice (2-3 months old) were measured under isoflurane anesthesia (initiation: 5% V/V in O_2_, maintenance: 1.5-2% V/V in O_2_), using 5 mg/kg carprofen (Rimadyl, Pfizer) and lidocaine (Braun) as analgetic.

The ultrafast Doppler setup consisted of an ultrasound research system (Vantage 256-LE, Verasonics, Redmond, WA, USA) interfaced with a high-frequency linear array (L22-14vX, Verasonics, operated at 15.6 MHz, aperture 12.8 mm). The ultrasound system was coupled to a workstation via a PCIe bridge to allow for fast (6 GB/s) and sustained data transfer. Data processing and Fourier domain image reconstruction was performed on a GPU (Titan GTX 2010, NVIDIA, Santa Clara, CA, USA) using custom-written C++/CUDA software. The setup was controlled using custom-written LabVIEW software (National Instruments, Austin, TX, USA).

With respect to the ultrasound data acquisition a pulse repetition frequency (determined by the time it takes to acquire the data for one ultrasound image) of 20 kHz was used and 20 angled (between -7 and + 7 degrees) planewaves were averaged for one compounded frame providing a Doppler pulse repetition frequency of 1 kHz. The compounded frames were streamed to disc for further offline processing. For every 2D vascular image, the Doppler signal of a set of 2270 compounded frames was used. Doppler processing consisted of SVD-based clutter filtering (Demené et al., 2015), followed by a 5^th^ order high-pass zero-phase Butterworth filter with a cut-off frequency of 60 Hz. Doppler frames with an average value of one standard deviation below or above the median value for all frames were discarded from the set as they are linked to high overall motion, for example induced by breathing. Spatial up sampling of the Doppler filtered frames to an isotropic grid of 25 µm was performed by zero padding in the frequency domain and the final ultrafast Doppler image was obtained by computing the power of the Doppler signal for every pixel. Log compression was applied to compress the visible dynamic range to a level of 25-35 dB depending on the noise-level.

### Transthoracic echocardiography

For echocardiography, mice were shaved and lightly anesthetized with 1.5% isoflurane. Non-invasive echocardiographic parameters were measured using a Vevo-2100 high frequency ultrasound system (VisualSonics Inc., Toronto, Canada) and analyzed using Vevo-2100-1.6.0 software. Long-axis EKG-triggered cine loops of the left ventricular (LV) contraction cycle were obtained in B-mode to assess end-diastolic/systolic volume. Short-axis recordings of the LV contraction cycle were taken in M-mode to assess wall thickness of the anterior/posterior wall at the mid-papillary level. From M-mode recordings, LV posterior wall thickness in systole (LV PWs) and diastole (LV PWd) were determined. LV mass was calculated as 0.8 * (1.04 * ((LVIDd + LV PWd + IVSd)^3) - (LVIDd)^3)) + 0.6 with LVid = LV internal dimension; fractional shortening (FS) was calculated as (LVIDd-LVIDs) / (LVIDd*100). Ejection fraction (EF) was calculated as 100 * (SV/Vd) with Vs = systolic volume (3.1416 * (LVIDs^3)/6), Vd = diastolic volume (3.1416 * (LVIDd^3)/6), and SV = stroke volume (Vd-Vs).

In vivo ultrasound imaging of the aortic arch, abdominal aorta and left ventricle (LV) was performed with a Vevo-2100 (Visualsonics Inc) using a 40-MHz linear interfaced array transducer (MS550S). B-mode and M-mode images of the aorta were captured. Diameters of the aortic arch were measured from the parasternal window at the level of the ascending aorta. Distensibility of the aortic arch was measured as the systolic to diastolic aortic diameter ratio in M-mode image data (calculated as systolic diameter minus diastolic diameter, divided by the diastolic diameter).

### MMP imaging

Per 25 grams of body weight 2 nmol specific MMP activatable NIRF probes, MMPSense680™ (Perkin Elmer Inc., Akron, Ohio, USA), was injected into the tail vein of anesthetized mice. Intact aortas and hearts were harvested 24 hours after injection for ex vivo fluorescence imaging and analyzed using the Odyssey Imaging system (LI-COR Biosciences, Lincoln, Nebraska, USA). Near-infrared images were obtained in the 800 nm channel.

### Cardiac histological analysis and (immunofluorescence) microscopy

Prior to histological analysis, hearts were perfused with 4% paraformaldehyde (PFA), followed by paraffin embedding and sectioning at 4 µm. These sections were cut and mounted on coated slides and dried overnight at 37C. Slides were dewaxed in xylene and hydrated using graded alcohols to tap water. The sections were subsequently stained with fluorescein isothiocyanate (FITC)-labeled wheat-germ-agglutinin (WGA; Sigma) to visualize and quantify the cell cross-sectional area; hematoxylin and eosin (H&E; Sigma) for routine histological analysis; and Sirius Red (Sigma) for detection of fibrillar collagen. For immunofluorescence, sections were dried, fixed with 4% PFA, and permeabilized with 0.1% Triton X-100 (Sigma), dissolved in 1% BSA in PBS for 10 min and blocked with 10% goat serum in PBS for 60 min. Then, the slides were incubated overnight with primary antibody diluted in 0.1% BSA in PBS at 4 °C. Secondary antibody incubation was performed at room temperature for 1 h, followed by 5 min Hoechst incubation and mounting with Fluoromount (Southern Biotech). Images were taken using an Olympus BX63 microscope and analyzed in ImageJ.

### Brain histological analysis

For the visualization of blood vessels in the brain, we used adult (>3 months) mice. Mice were anesthetized with pentobarbital (80 mg/kg i.p.) and transcardially perfused first with 4% paraformaldehyde and then with 10% Indian ink mixed with 2% gelatin (preheated to 40 °C), based on a previously published method (Xue et al., 2014). After injection of the ink, the mice were placed with their ventral side up for 10-20 min at -20 °C to solidify the gelatin. Afterwards, the brains were extracted, fixed for 2 h in 4% paraformaldehyde, and incubated overnight in 0.1 M phosphate buffer (PB) with 10% sucrose at 4 °C. Next, the brains were embedded in 14% gelatin/10% sucrose and postfixed for 2 h in 10% formaldehyde/30% sucrose. After another overnight incubation, this time in 0.1 M PB with 30% sucrose, 40 µm sagittal sections were cut on a sliding microtome while being cooled with dry ice. The sections were processed freely floating, washed twice with 0.1 M phosphate-buffered saline (PBS), blocked for endogenous peroxidase activity (1% H_2_O_2_ in 0.1 M PBS), washed again twice in PBS, and were stained for 3 minutes in a 1 % thionine solution, followed by a differentiation of the sections in 96% ethanol, followed by dehydration using several steps with 100% ethanol and xylene, and cover slipped with Permount mounting medium (Fisher Scientific, Waltham, MA, USA).

In the section approx. 100 µm lateral of the midline, areas of 650×750 µm were chosen for counting the number of loops under a light microscope. For each mouse, we counted the loops in the hippocampus, cerebellar cortex and brainstem. The total number of loops per brain region, per mouse was reported.

Sections from the same animals were used for immunofluorescent staining of GFAP, Iba1 and Fibulin 4 as previously described (Birkisdóttir et al., 2022), with chicken anti-GFAP (Abcam ab4674, 1∶5000), rabbit anti-Iba1 (WAKO Chemicals, 1∶ 2000) and rabbit anti-Fibulin 4 (1:100, provided by T. Sasaki (Sasaki et al., 2016)) as primary antibodies, and Alexa fluor 488 and cyanine 3 (Cy3)-conjugated as secondary antibodies raised in donkey (Jackson ImmunoResearch, West Grove, PA). Sections were stained with DAPI to visualize cell nuclei, mounted on coverslips, and placed on glass slides with Mowiol mounting medium, and imaged using a ZEISS Axio Imager M2 fluorescence microscope with Plan-Apochromat 10× objective and ZEN software for image acquisition and stitching. In addition, sections were imaged with a Zeiss LSM700 confocal laser scanning microscope with a Plan-Apochromat 63x/1.40 Oil objective.

In sagittal slices, we traced the Purkinje cell layer of each lobule in the cerebellar vermis, hemispheres and (para)flocculus, and measured its length using Neurolucida (MBF Bioscience, Williston, VT, USA). Typically, we analyzed approximately six slices for the vermis, twelve for the hemispheres, four for the flocculus, and eight for the paraflocculus. Vermal lobule X was visible in less slices, and lobule I was often missing. For each lobule, we took the largest perimeter found for each mouse. Overview pictures were made using a brightfield microscope (Nanozoomer 2.0-RS, Hamamatsu). Rostrocaudal and ventrodorsal diameters of the inferior colliculus were measured in the sagittal section with the largest inferior colliculus for each mouse.

### Statistical analyses

When the data were not normally distributed, non-parametric tests were performed. When applicable, corrections for multiple comparisons were made as indicated in the text. Differences were considered significant when *P* <0.05.

## Acknowledgments

Financial support for this study was provided by Health Holland to promote public-private partnerships (TKI-LSH EMCLSH21017; L.W.J.B.; EMCLSH20024, H.elA.), ZonMw Veni (91619109; L.K.), Erasmus Medical Center Fellowship (L.K.), the Medical Delta Programme (L.K. & C.I.D.Z), LSH-NWO Crossover (INTENSE; L.K. & C.I.D.Z.), NWO-ENW (OCENW.M.21.218; M.C.S.) and DBI2 of the NWO-Gravitation Program (C.I.D.Z.). This publication is part of the project CUBE (with project number 108845) of the research programme NWO-Groot, which is partly financed by the Dutch Research Council (NWO; B.S.G., S.K.E.K., P.K., C.I.D.Z.).

