## Supplemental information for "A long-deletion mouse model of Williams syndrome reveals *Ncf1*-dependent modulation of vascular and neural phenotypes"

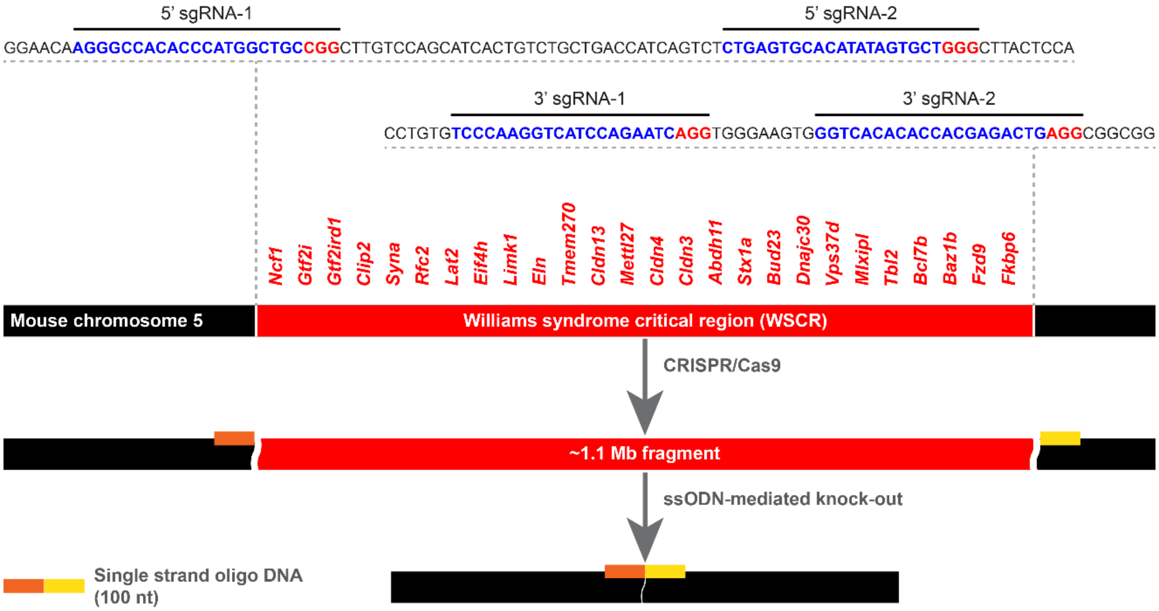

**Figure S1. CRISPR/Cas9-mediated excision of the Williams syndrome critical region (WSCR).** The LD mouse model was created by cutting out an approximately 1.1 Mb fragment of mouse chromosome 5 using CRISPR/Cas9-mediated excision.

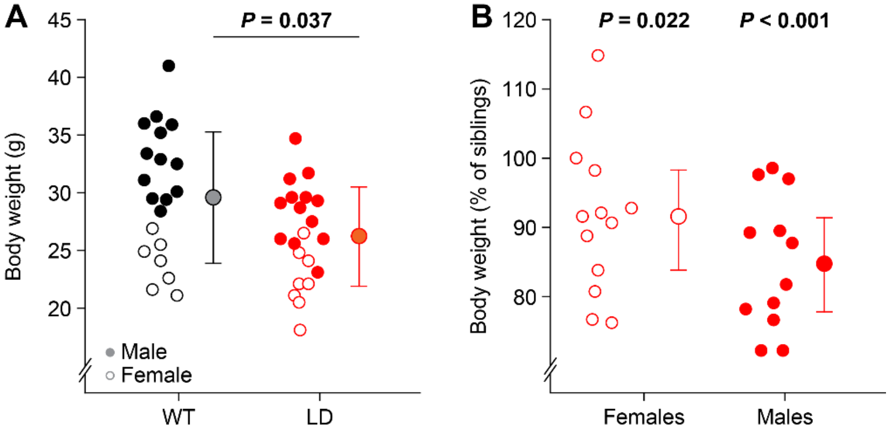

**Figure S2. Reduced body weight in LD mice.**
(*A*) Adult LD mice (6-8 months of age) had a reduced body weight compared to WT littermates ($n$ = 21 mice/genotype). $P$ = 0.037, $t$ = 2.168, $t$ test. (*B*) Body weights of female and male LD mice expressed as percentage of the average body weight of their WT sisters and brothers, respectively (>3 months of age, $n$ = 13 female and $n$ = 12 male LD mice). Females: $P$ = 0.022, $t$ = 2.633; males: $P$ < 0.001, $t$ = 5.403, $t$ tests, both are significant after Bonferroni correction. Averages ± sd.

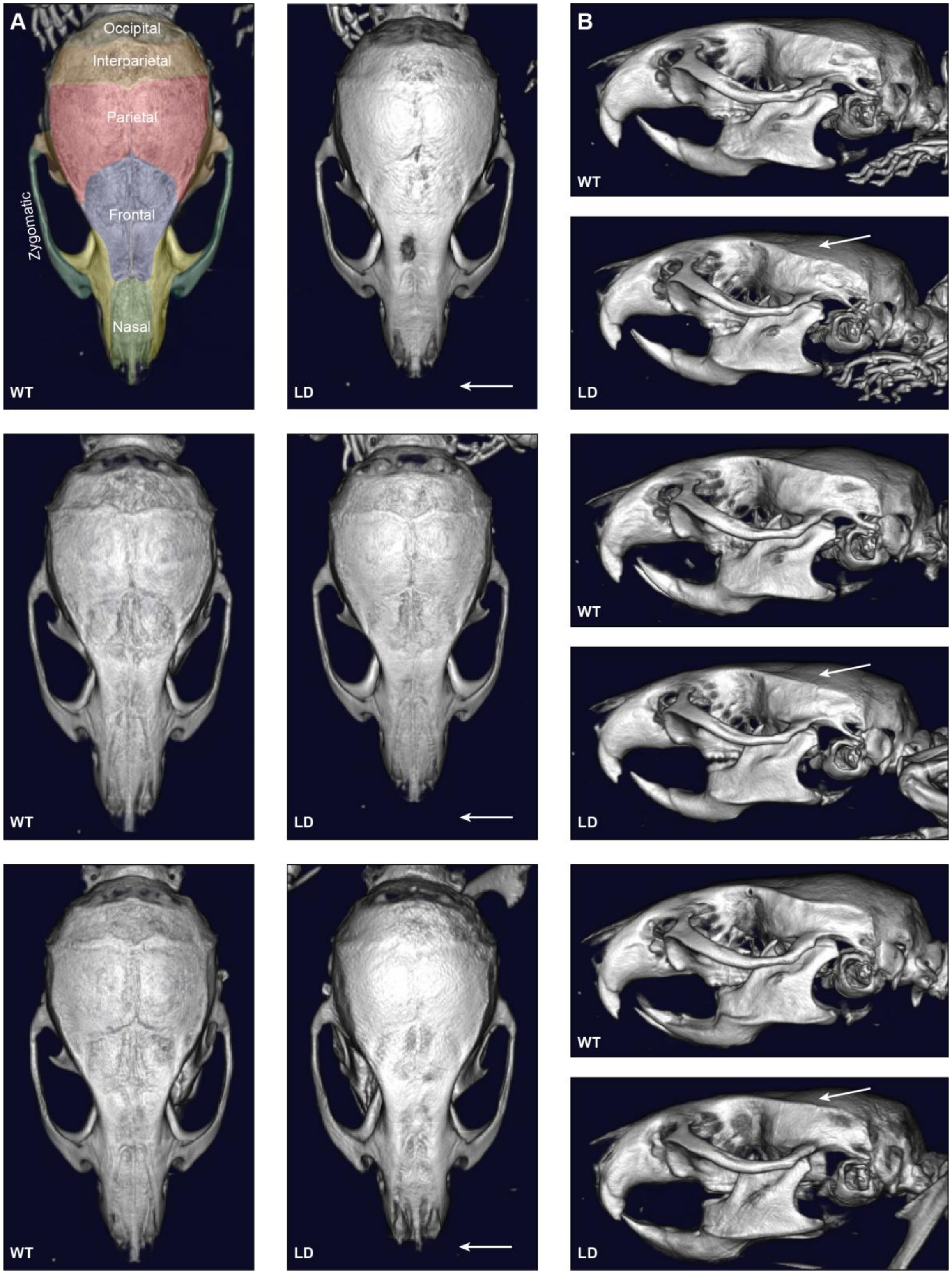

**Figure S3. Cranial abnormalities in LD mice.**
*(A)* Dorsal view of adult male mouse skulls generated with µCT scans. Every row depicts a pair of two littermates. LD mice have a shorter nasal bone (arrows). The middle row depicts the same images as Fig. 1C. *(B)* Side views. The arrows point towards a flatter bone structure in LD mice.

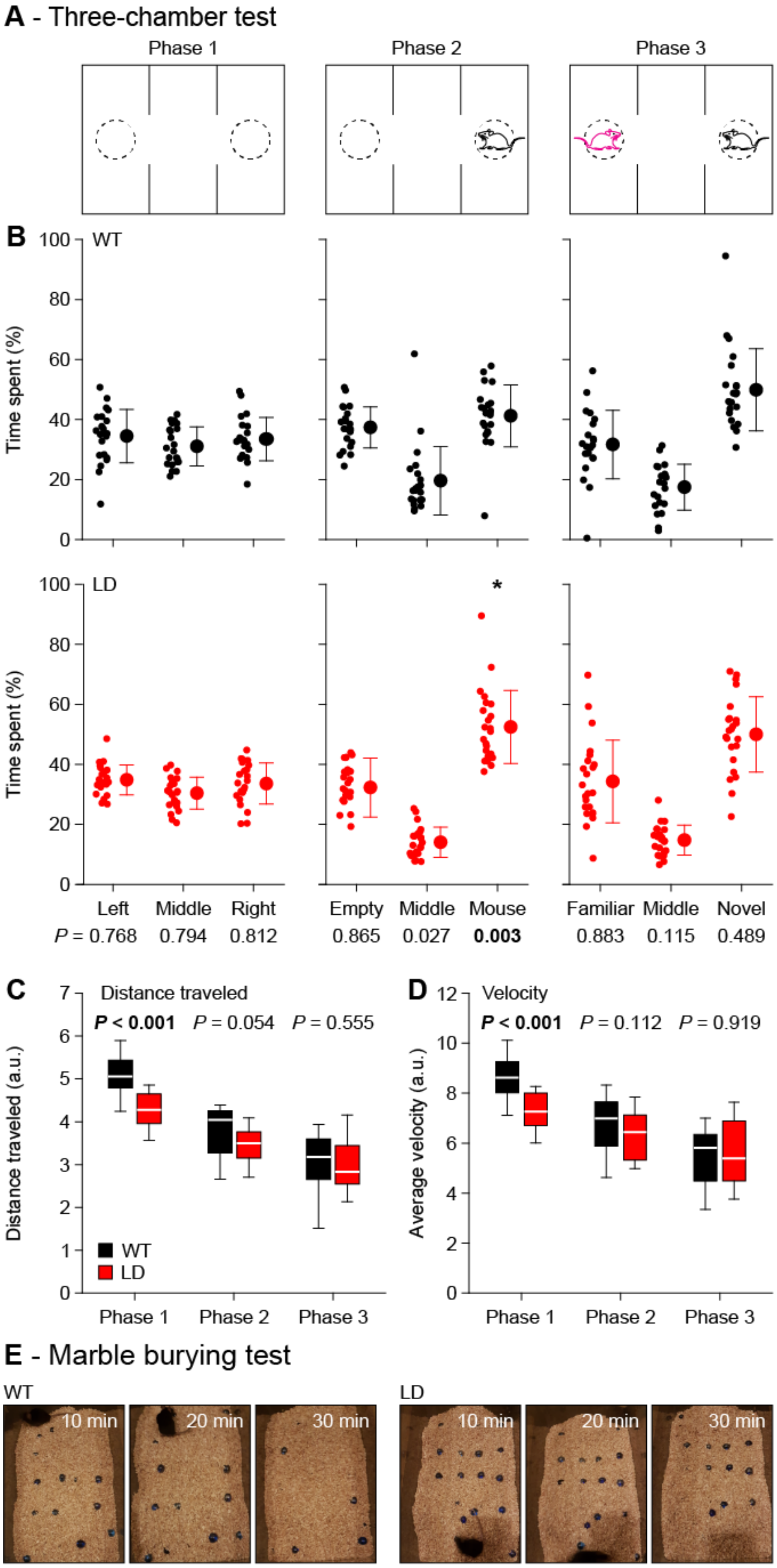

**Figure S4. Hyper-social and less compulsive behavior in LD mice.** (*A*) Three-chamber test phase 1: the test mouse can explore the environment freely. Phase 2: another mouse is present in a confinement in an outer chamber. Phase 3: a second, non-familiar mouse (red) is also present. (*B*) Only time spent in the chamber with the other mouse during phase 2 differed significantly between WT and LD mice ($n$ = 22 WT and 23 LD mice, $P$ = 0.003, Mann-Whitney test). (*C*) LD mice traveled less than WT mice during the first, but not during later phases ($n$ = 22 WT and 23 LD mice, $P$ < 0.001, Mann-Whitney test). (*D*) The same for velocity ($n$ = 22 WT and 23 LD mice, $P$ < 0.001, Mann-Whitney test). Significance is indicated after Benjamini-Hochberg correction for multiple comparisons. Population statistics indicated as averages ± sd. (*E*) Pictures at 10, 20 and 30 min of marble burying.

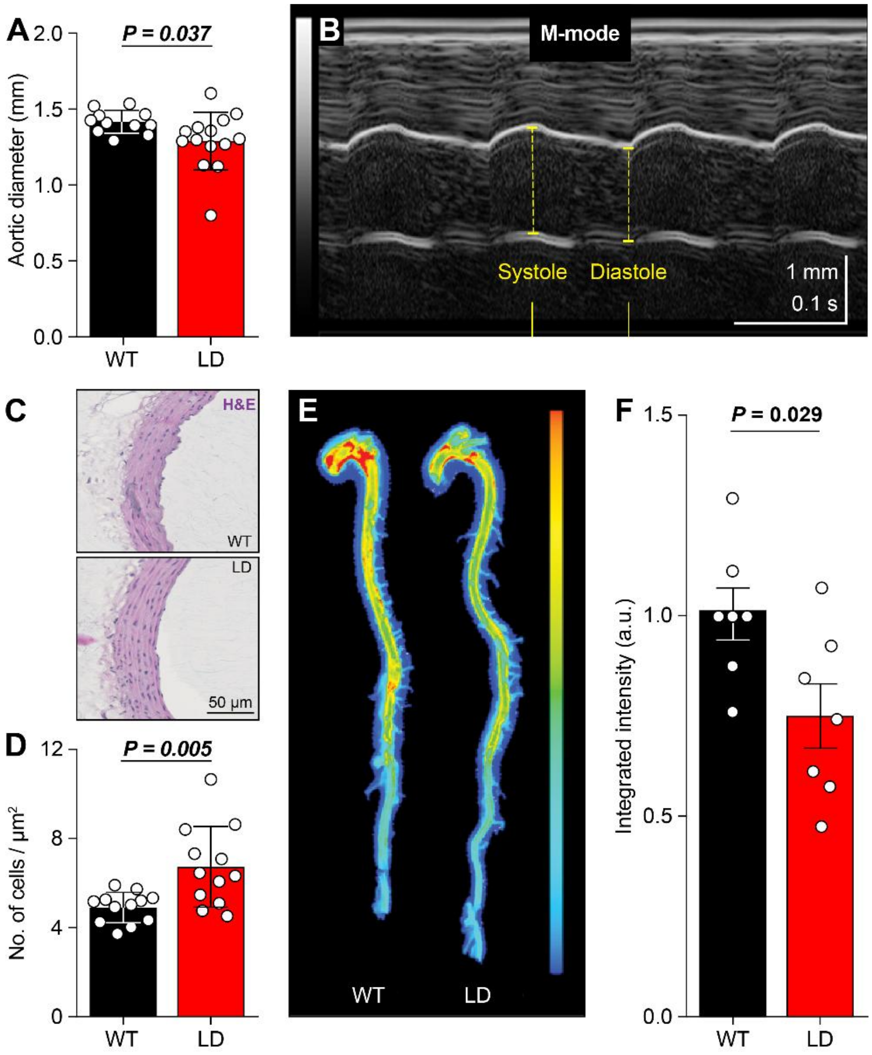

**Figure S5**. **LD mice have a stiffer aorta with a thicker, but stable wall.** (*A*) The ascending aorta has a narrower lumen in LD mice ($n$ = 11 WT and 14 LD mice, $P$ = 0.037, $t$ = 2.261, $t$ test). (*B*) Functional ultrasound imaging of a representative mouse showing the characterization of aortic distensibility, which is the difference between the diameter during systole vs. diastole (dashed yellow lines). See also Fig. 2C-D. (*C*) Hematoxylin and eosin (H&E) staining demonstrating the increased number of cells in the aortic wall of LD mice (*D*, $n$ = 12 mice/genotype, $P$ = 0.005 Mann-Whitney test). (*E*) Representative scans showing decreased levels of matrix metalloprotein (MMP) activity in the aortas of LD mice. (*F*) Fluorescence due to MMP activity was actually lower in LD mice (n = 7 mice/genotype, $P$ = 0.029, $t$ = 2.514, $t$ test). Population statistics depicted as averages ± sd.

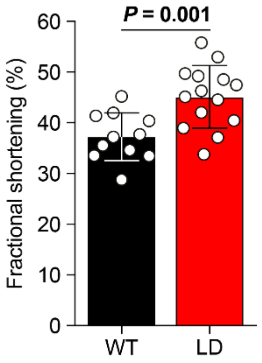

**Figure S6. Decreased elasticity of the aorta.**
Increased fractional shortening (*n* = 11 WT and 14 LD mice, *P* = 0.002, Mann-Whitney test). Averages ± sd.

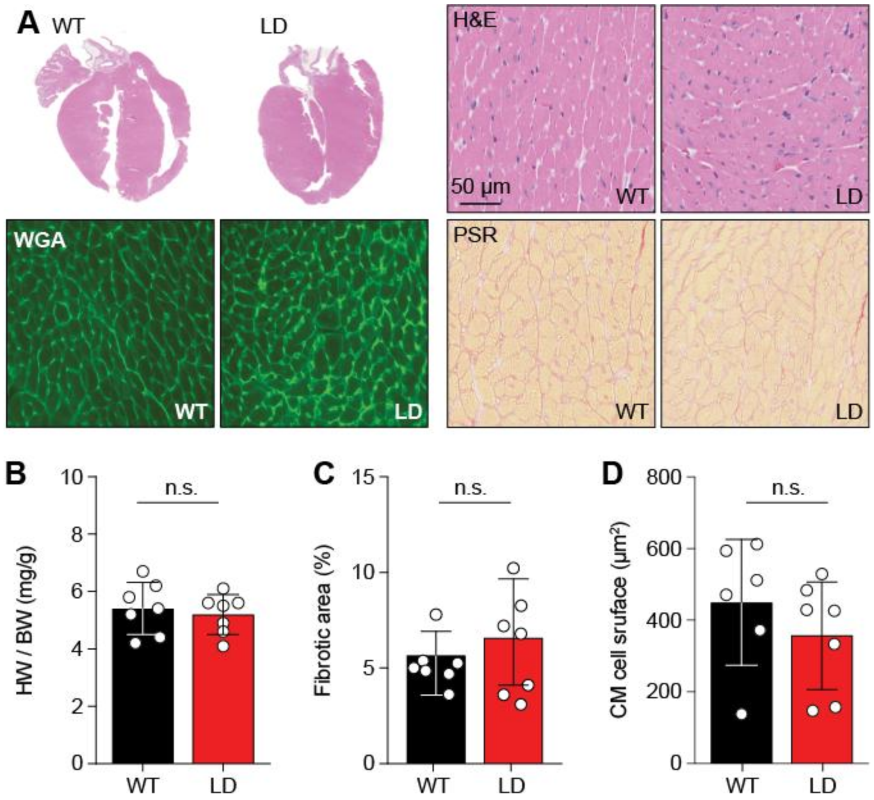

**Figure S7. Unaltered cardiac anatomy.**
(*A*) Representative examples of whole hearts and cardiac sections with H&E, wheat germ agglutinin (WGA) and picrosirius (PSR) stainings, all showing relatively normal cardiac anatomy. (*B*) Gravimetric analysis of corrected heart weights (*n* = 7 mice/genotype, *P* = 0.686, Mann-Whitney test). (*C*) Quantification of collagen density in cardiac tissue sections. (*n* = 7 mice/genotype, *P* = 0.068, Mann-Whitney test). (*D*) Analysis of cardiomyocyte (CM) hypertrophy in cardiac tissue sections. Slices (6 µm) of the left ventricular myocardium were stained with Alexa 647–labeled WGA for determination of myocyte cross-sectional areas. (*n* = 6 WT and 7 LD mice, *P* = 0.366, Mann-Whitney test). Averages ± sd.

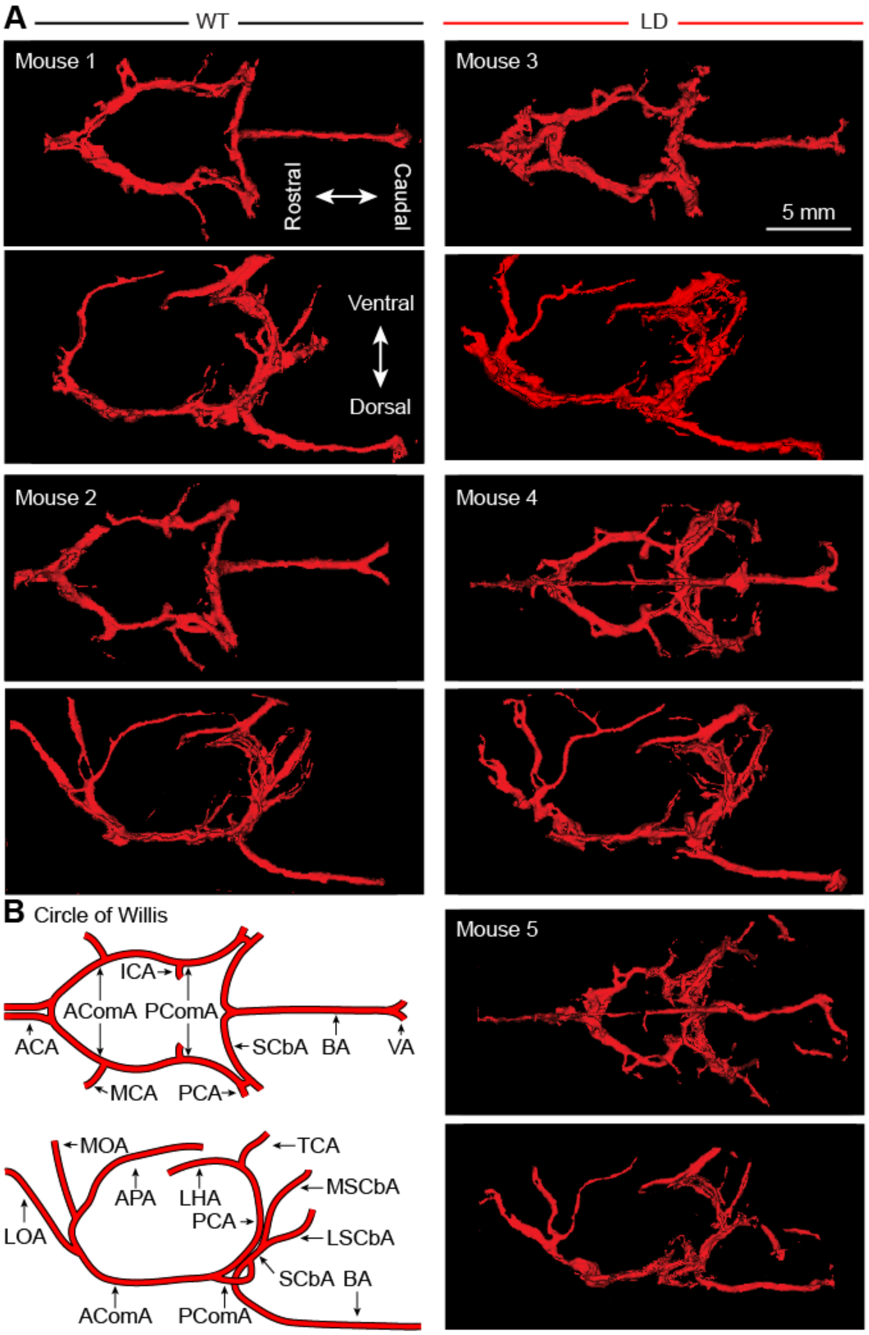

**Figure S8. Altered circle of Willis in LD mice.**

(*A*) Intracranial vasculature of five mice visualized using µCT scans. (*B*) Schematic representation of the different arteries that can be identified in the images shown in *A*. ACA = anterior cerebral artery (a.), AComA = anterior communicating a., APA = azygos pericallosal a., BA = basilar a., ICA = internal carotid a., LHA = longitudinal hippocampal a., LOA = lateral orbitofrontal a., LSCbA = lateral superior cerebellar a., MOA = medial orbitofrontal a., MSCbA, medial superior cerebellar a., PCbA = posterior cerebellar a., PComA = posterior commissural a., SCbA = superior cerebellar a., TCA = transverse collicular a., VA = vertebral a.

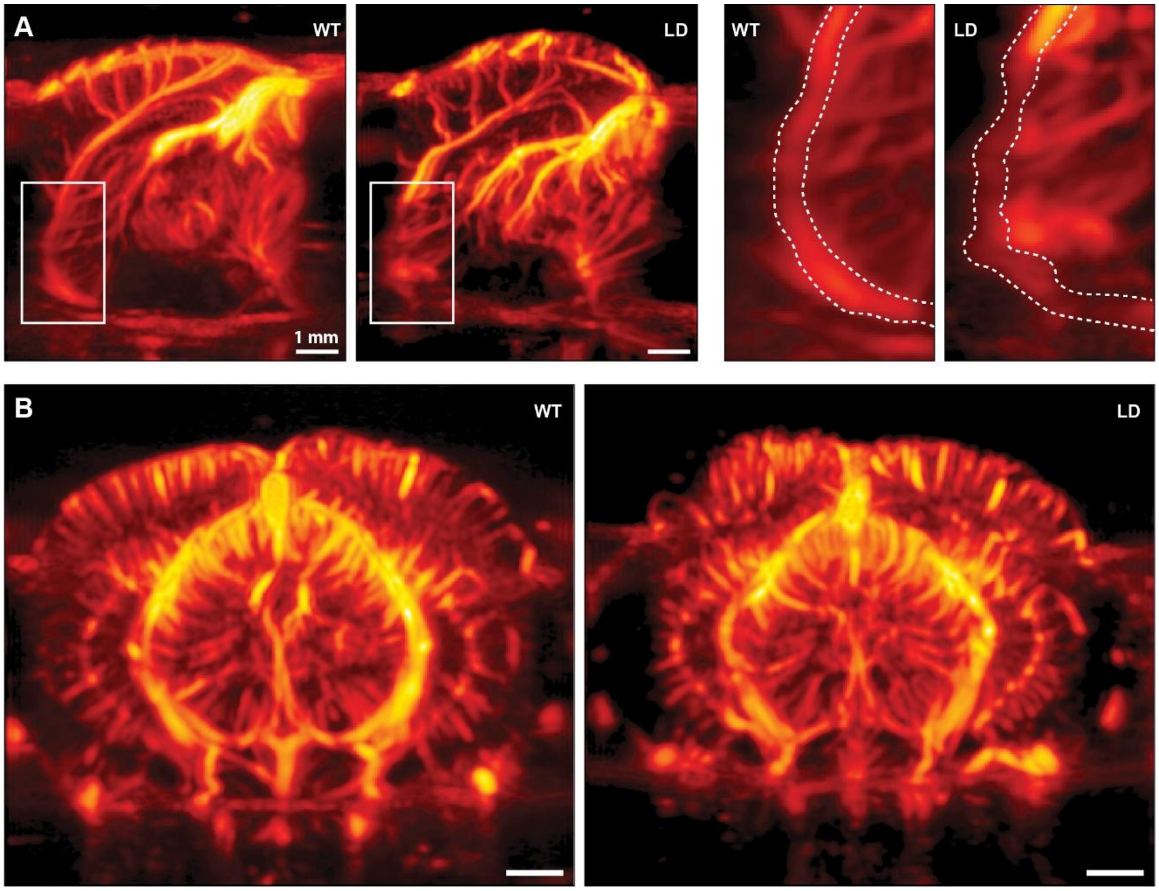

**Figure S9. Altered brain vasculature in LD mice.**
(*A*) Sagittal view of the brain vasculature in WT and LD mice visualized using in vivo ultrafast Doppler imaging of two representative mice. White rectangles indicate areas enlarged in the right columns. (*B*) Coronal view.

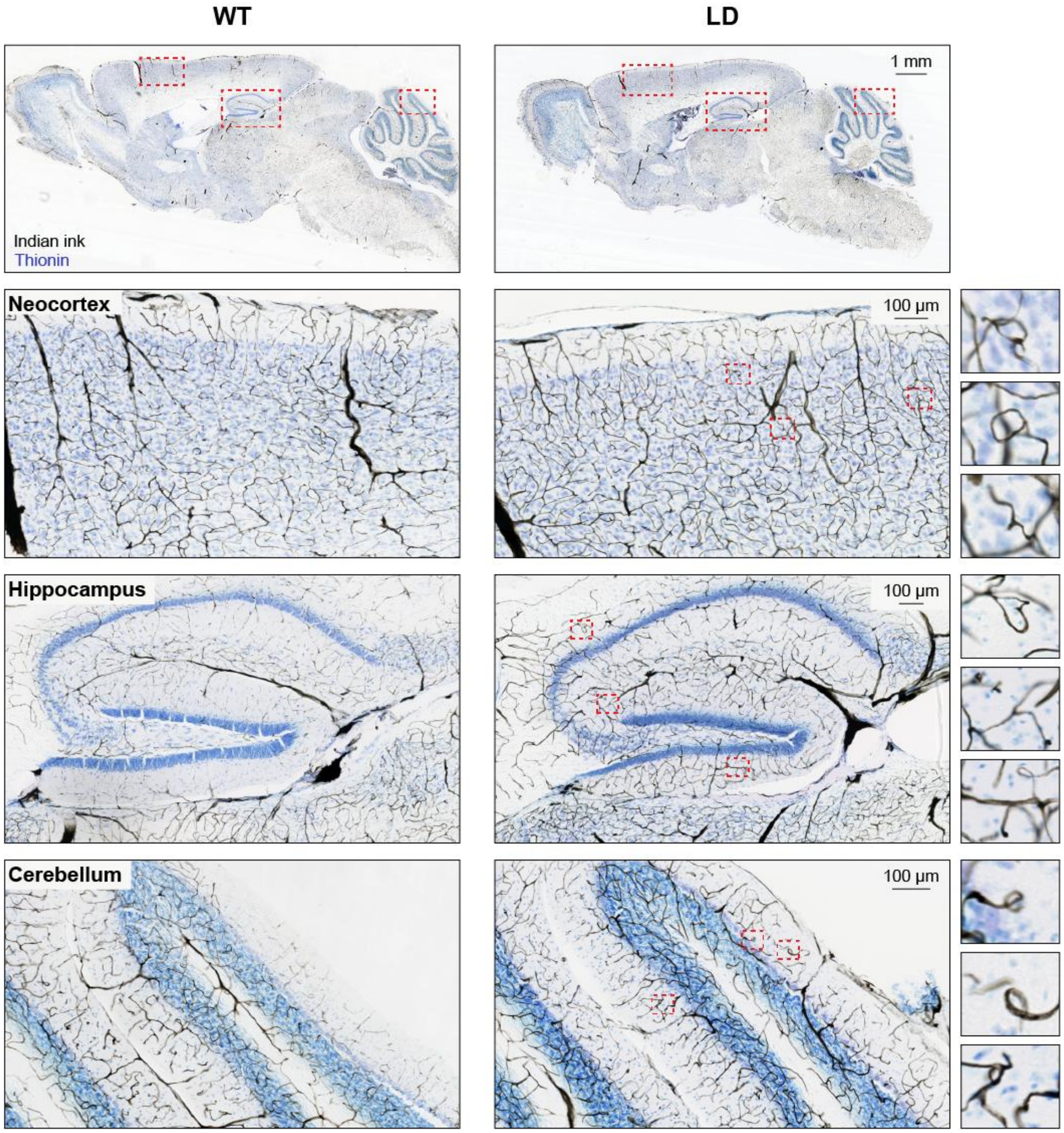

**Figure S10. More irregular elements in the microvasculature of LD mouse brains.**
Irregular elements in the blood vessels stained with Indian ink in 50 µm thick brain slices can be found in different regions, and include hairpins and spirals.

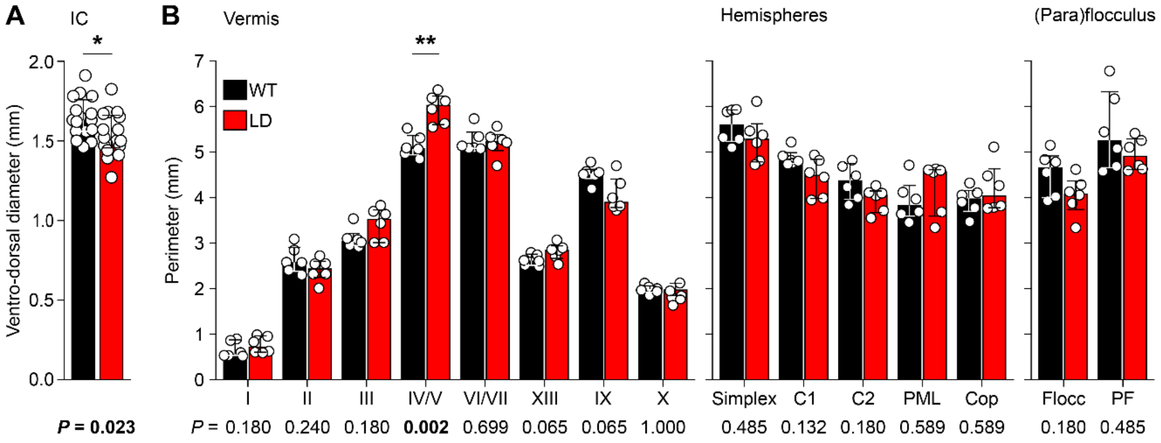

**Figure S11. Specific alterations in regional brain volumes in LD mice.**
(*A*) LD mice had a smaller inferior colliculus as measured along the ventro-dorsal axis ($n$ = 16 WT and 17 LD mice, $P$ = 0.023, $t$ = 2.395, $t$ test). See Fig. 5A-B. (*B*) The perimeter (see Fig. 5A) of the vermal lobules were similar between WT and LD mice, with exception of lobule IV/V that was enlarged in LD mice (see Fig. 5C; $n$ = 6 mice/genotype, Mann-Whitney tests, ** significant after Bonferroni correction for multiple comparisons). (*C*) The lobules of the hemispheres had similar perimeters in WT and LD mice ($n$ = 6 mice/genotype, Mann-Whitney tests). (*D*) Also the flocculus and the paraflocculus were relatively normal in LD mice ($n$ = 6 mice/genotype, Mann-Whitney tests). Averages ± sd.

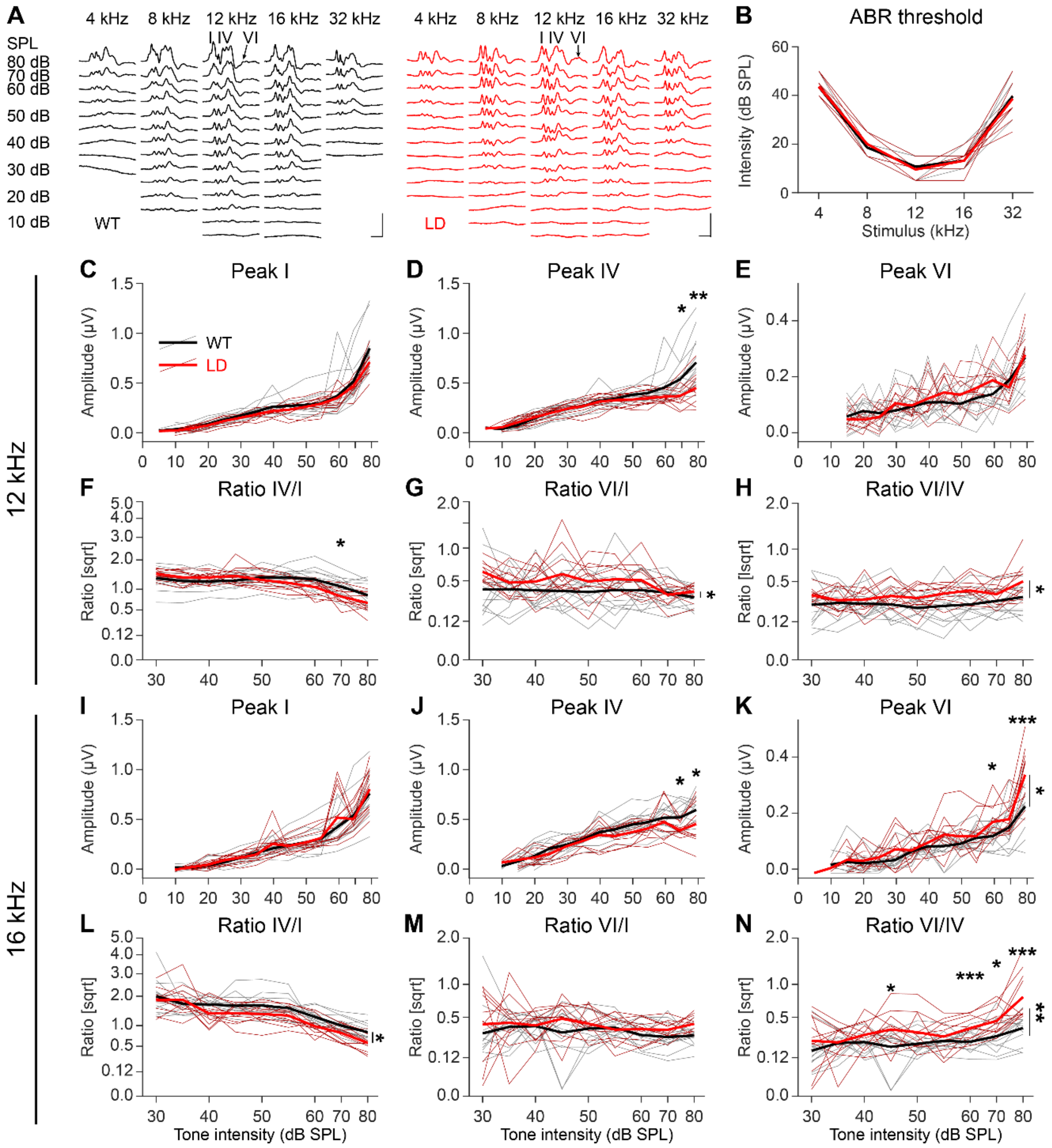

**Figure S12. Normal hearing threshold, but reduced responses to loud sounds in LD mice.** (*A*) Auditory brainstem responses (ABR) evoked by 1 ms pure tone pips of different frequencies and intensities. Traces are averages from the two lead electrodes with an inverted sign. Scale bars: 1µV and 2 ms. (*B*) ABR thresholds for different frequencies. Amplitudes of peaks I (*C*), IV (*D*) and VI (*E*) at different sound intensities for a 12 kHz tone. Amplitude ratios of peaks IV/I (*F*), VI/I (*G*) and VI/IV (*H*). (*I*-*N*) The same for a 16 kHz tones. Thin lines indicate individual mice, thick lines the population averages. Statistical evaluation in Tables S2 and S3. * $P < 0.05$, ** $P < 0.01$, *** $P < 0.001$.

Balance beam

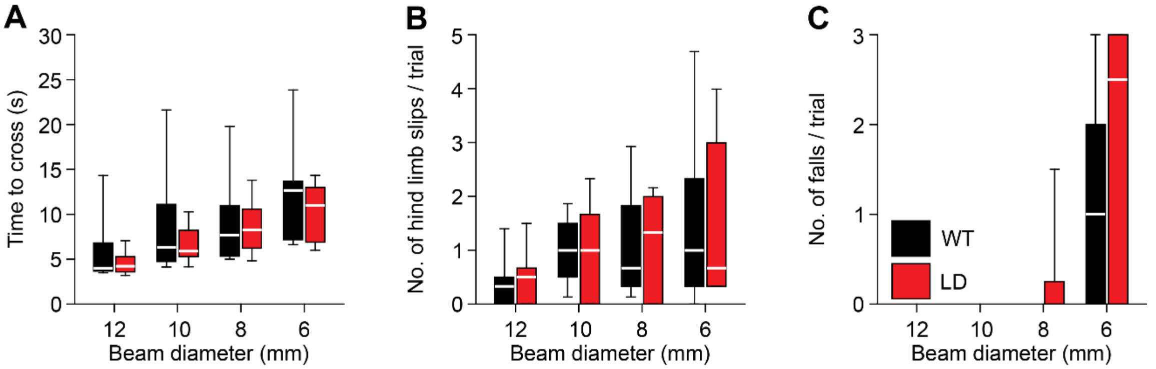

**Figure S13. Apart from motor coordination, normal performance on the balance beam.** (*A*) Time to cross the balance beam was unaffected by the LD mutation. Box plots of median and IQR, with the whiskers indicating the $10^{th}$ and $90^{th}$ percentiles, of the averages of three replicates per mouse per beam diameter. Only completed trials were included in this analysis ($n$ = 14 WT and 13 LD mice, 12 mm: $P$ = 0.627, 10 mm: $P$ = 0.680, 8 mm: $P$ = 0.789, 6 mm: $P$ = 0.618, Mann-Whitney tests). (*B*) Also the occurrence of hindlimb slips was not increased in LD mice. Only completed trials were included in this analysis (12 mm: $P$ = 0.364, 10 mm: $P$ = 0.941, 8 mm: $P$ = 0.864, 6 mm: $P$ = 0.927, Mann-Whitney tests). (*C*) Number of trials in which the mice did not reach the other side of the beam (6 mm: $P$ = 0.285, Mann-Whitney test).

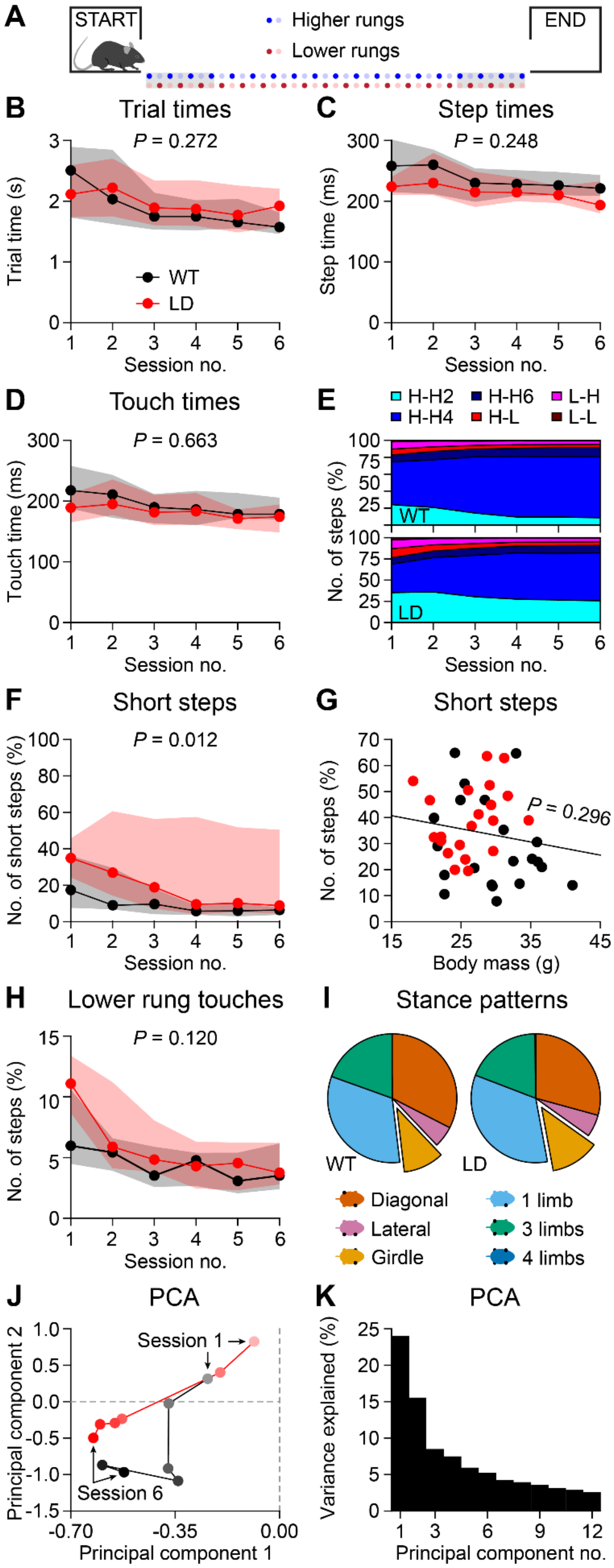

**Figure S14. Relatively normal locomotor performance on the ErasmusLadder.**

(*A*) The ErasmusLadder consists of a horizontal ladder with alternating higher and lower rungs between two shelter boxes. (*B*) Trial times were not affected by the LD mutation ($n$ = 21 mice per genotype, $P$ = 0.272, $F$ = 1.240, repeated measures ANOVA). (*C*) Step times ($P$ = 0.248, $F$ = 1.375, repeated measures ANOVA). (*D*) Duration of rung touches ($P$ = 0.663, $F$ = 0.193, repeated measures ANOVA). (*E*) Distribution of step types: the letters indicate the rung types (H = higher, L = lower) and the number the step size. Thus, H-H2 is a step from a higher rung to the next higher rung (skipping the lower rung in between). (*F*) The relative occurrence of short steps (H-H2) was higher in LD mice ($P$ = 0.012, $F$ = 6.913, repeated measures ANOVA). (*G*) Given the large variability between mice in the occurrence of short steps, and a putative interaction with body weight, we made a linear regression between body weight and the percentage of short steps during session 1 (R = 0.165, $P$ = 0.296, Pearson's correlation). (*H*) Overall, LD mice did not have a tendency to touch lower rungs more often than WT littermates ($P$ = 0.120, $F$ = 2.529, repeated measures ANOVA). (*I*) Distribution of the step types. See also Fig. 5F. (*J*) Principal component analysis (PCA) revealed a similar, but not identical pattern for both genotypes. (*K*) Variance explained for each principal component.

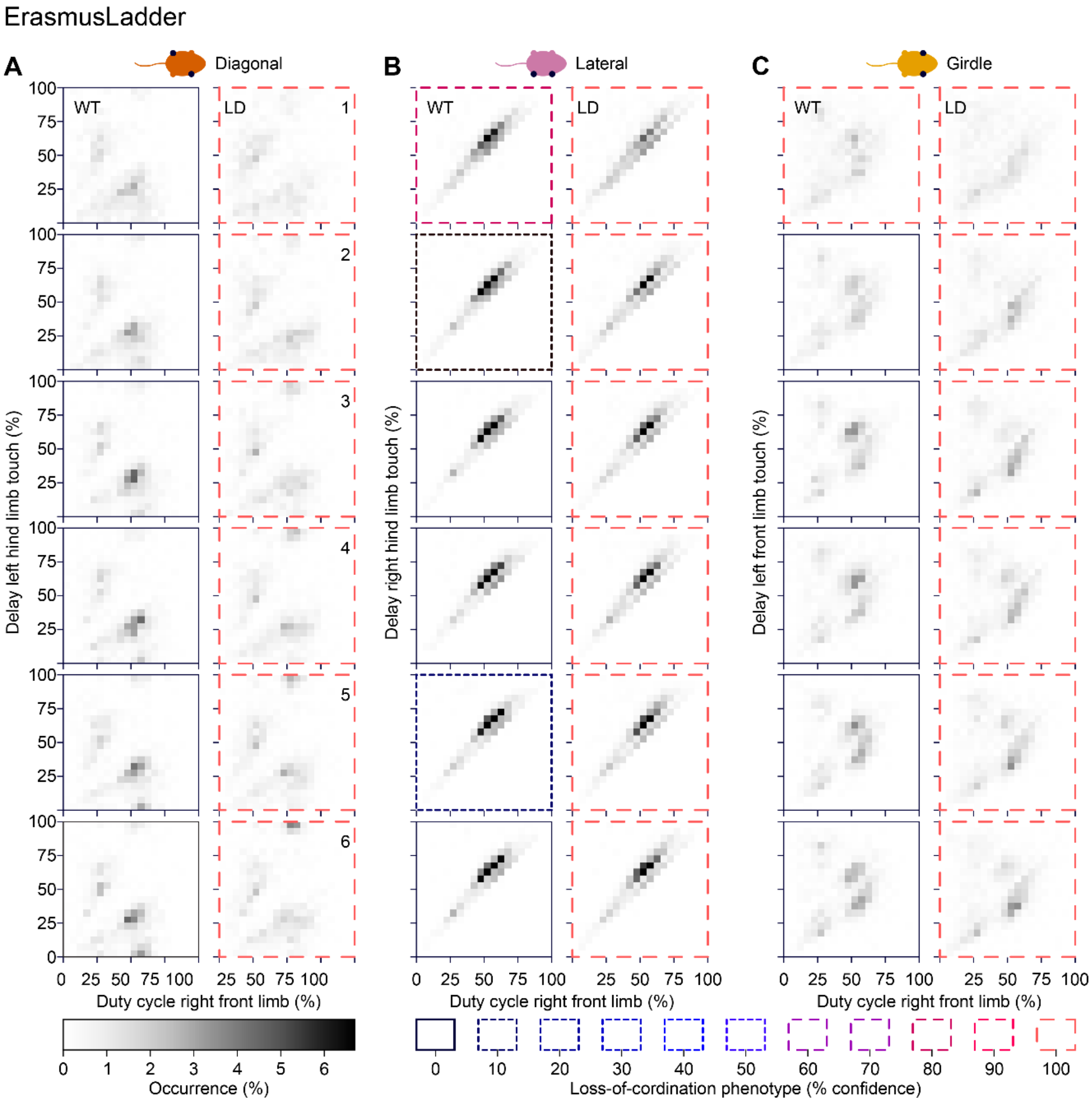

**Figure S15. LD mice show impaired inter-limb coordination.**
(*A*) Hildebrand plots representing the combinations of duty cycle (= percentage of step cycle during which the right forelimb touches a rung) and the delay in the step cycle between placement of the right forelimb and the left hindlimb. All registered steps from all mice were grouped per session and genotype. The same for the comparison between the duty cycle from the right frond limb and the right hindlimb (*B*) or the left forelimb (*C*). The thick colored borders indicate the sessions that were marked as deviant by unsupervised machine learning, showing that LD mice were discernible from WT littermates with respect to their coordination over all three body axes.

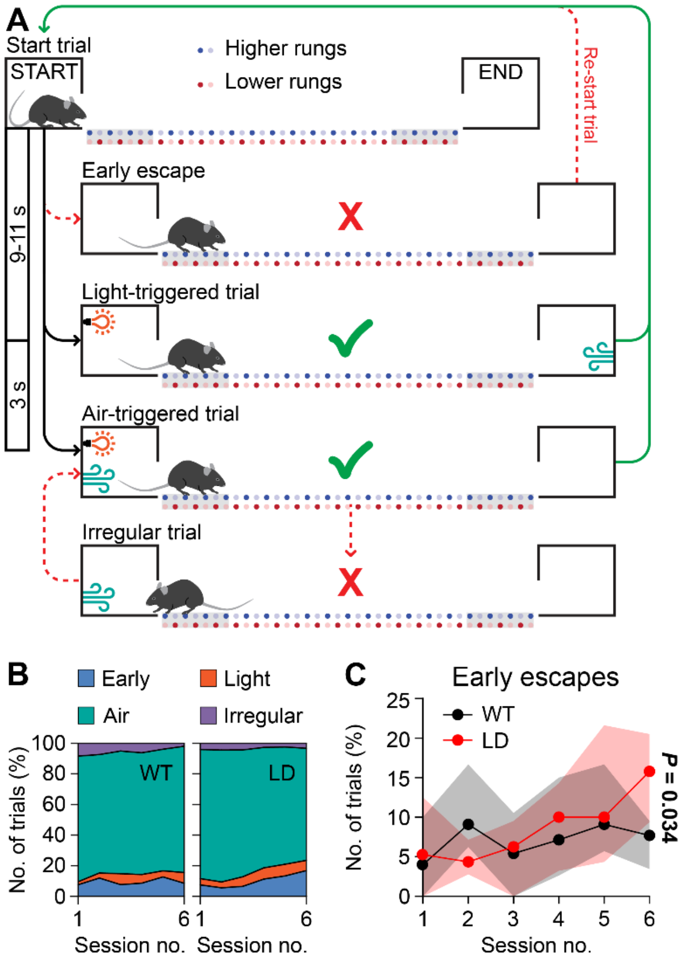

**Figure S16. LD mice did not start the trials on the ErasmusLadder earlier or later.**
(*A*) Temporal structure of the trials. The start of a trial was announced by lighting an LED in the start box. Early escapes, i.e., departures from the start box before the LED was turned on, were discouraged by a strong head-wind blowing over the ErasmusLadder. Early escapes induced the abortion of a trial, and that trial was then repeated until the mice left after lighting the LED. Data form early escapes were not included in the analysis of gaiting. Three seconds after the LED was turned on, air flow in the start box started. Once the mouse was on the rungs, a strong tail-wind encouraged the mouse to walk efficiently to the end box. Returns to the start box occasionally occurred, and the steps made during these unfinished crossings were excluded from the analysis of gaiting. 9-11 seconds after arriving in the end box, the next trial started, during which the mice crossed the ErasmusLadder in the opposite direction. (*B*) Distribution of the start moments. (*C*) LD mice started their trials earlier (before the LED was turned on) during the later sessions (genotype x time interaction: $P$ = 0.034, $F$ = 2.918, df = 3.204, repeated measures ANOVA with Greenhouse-Geisser correction). Panel *A* is reproduced with permission from (Jaarsma et al., 2024).

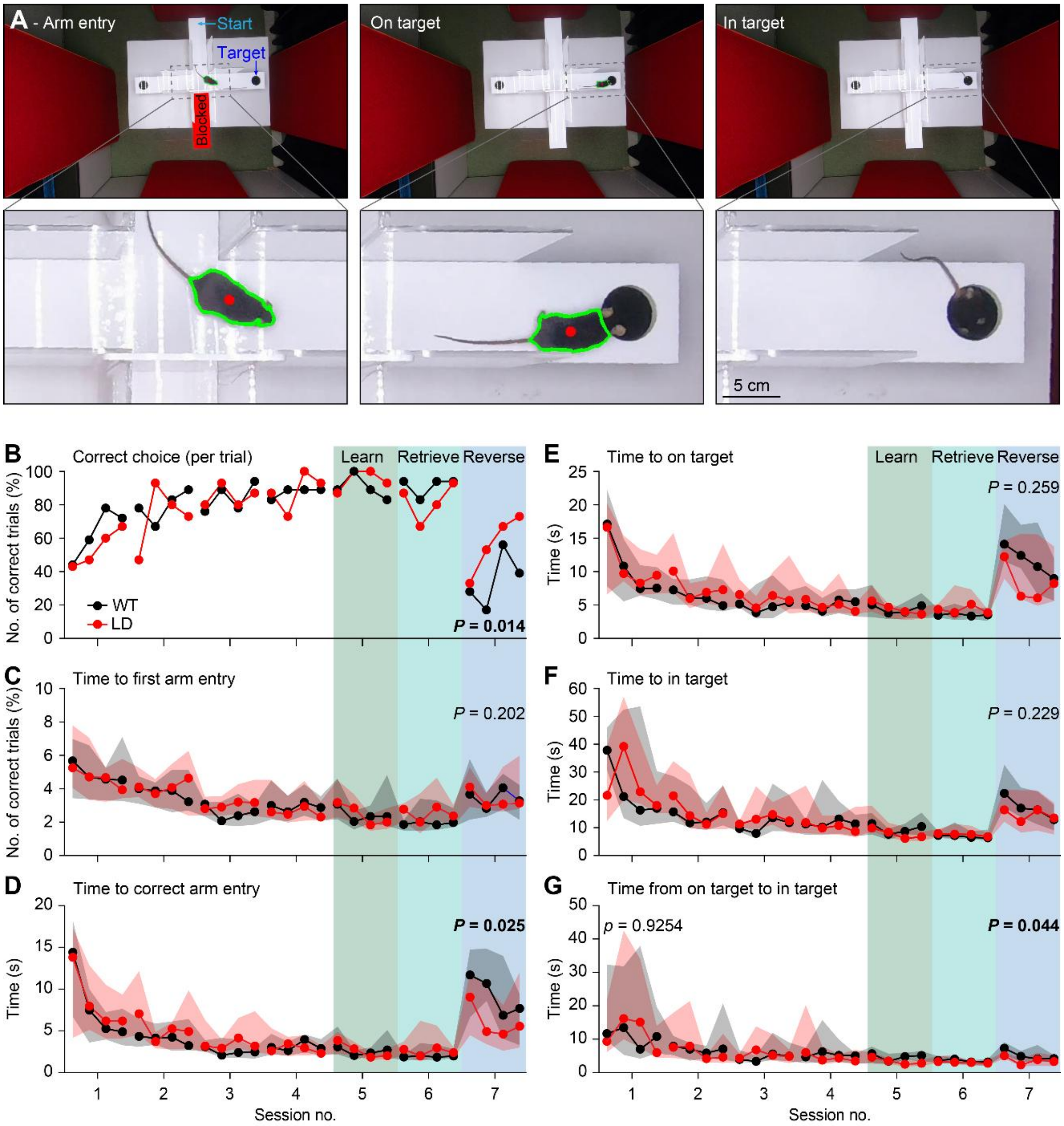

**Figure S17. LD mice have no problems learning to find the target on a T-maze, and are faster in reversal learning than WT littermates.**

(*A*) Mice start on the middle arm of a T-maze and could escape to a shelter underneath one of the other arms. After 5 days with 4 trials each, session 6 was 1 week later and during session 7, the mice started from the opposite arm, so that they had to change their trajectory. (*B*) Correct choices for first arm entry per trial. Since only the last session (reversal) showed a difference between WT and LD mice (see Fig. 5G), we performed statistical analysis only for that session here ($n$ = 18 WT and 15 LD mice, $P$ = 0.014, Mann-Whitney test). (*C*) Time to first entry ($P$ = 0.202, Mann-Whitney test). (*D*) Time to correct arm entry ($P$ = 0.025, Mann-Whitney test). (*E*) Time to target area (both ears should be in the target) ($P$ = 0.259, Mann-Whitney test). (*F*) Time to escape in target (all four limbs should be in the target) ($P$ = 0.229, Mann-Whitney test). (*G*) Time from reaching the target until escaping in the target ($P$ = 0.044, Mann-Whitney test).

**Table S1. Previously published mouse models**

| | Complete deletion (CD) | Proximal deletion (PD) | Distal deletion (DD) | D/P deletion |
|---|---|---|---|---|
| Background | C57BL6 | ~85% C57BL/6J and ~15% 129SvEv. | ~85% C57BL/6J and ~15% 129SvEv. | ~85% C57BL/6J and ~15% 129SvEv. |
| Deletion | *Gtf2i – Fkbp6* (Segura-Puimedon et al., 2014) | *Gtf2i – Limk1* (Li et al., 2009) | *Limk1 – Fkbp6* (Li et al., 2009) | *Gtf2i – Fkbp6*, with *Limk1* HOM KO (Li et al., 2009) |
| Vital parameters | | | | |
| Weight (at birth) | Normal (Segura-Puimedon et al., 2014; Giannoccaro et al., 2023) | | | |
| Weight (development) | Reduced (Giannoccaro et al., 2023) | | | |
| Weight (adult) | Reduced (Segura-Puimedon et al., 2014; Jiménez-Altayó et al., 2020) | Reduced (Li et al., 2009) | Reduced (Li et al., 2009) | Reduced (Li et al., 2009) |
| Lifespan | Normal (Segura-Puimedon et al., 2014) | Normal (Li et al., 2009) | Normal (Li et al., 2009) | Normal (Li et al., 2009) |
| Cardiovascular phenotype | | | | |
| Arterial pressure | Increased (Segura-Puimedon et al., 2014) | | | |
| Systolic blood pressure | Normal (Jiménez-Altayó et al., 2020) | | Increased (Goergen et al., 2011) | Normal (Goergen et al., 2011) |
| Heart rate | Normal (Segura-Puimedon et al., 2014) | | | |
| Heart size | Increased (Segura-Puimedon et al., 2014) | | | |

| | | | | |
|---|---|---|---|---|
| Aortic arch | Elongated (Jiménez-Altayó et al., 2020) | | | |
| Elastic lamellae (aorta) | Thinner (Jiménez-Altayó et al., 2020) | Normal (Goergen et al., 2011) | Disorganised (Goergen et al., 2011) | Disorganised (Goergen et al., 2011) |
| Elastin layers (aorta) | Increased (Jiménez-Altayó et al., 2020) | Normal (Goergen et al., 2011) | Normal (Goergen et al., 2011) | Normal (Goergen et al., 2011) |
| Wall thickness (aorta) | Normal (Jiménez-Altayó et al., 2020) | | | |
| Lumen (ascending aorta) | Smaller (Jiménez-Altayó et al., 2020) | | | |
| ACh-induced vasodilatation (aorta) | Normal (Jiménez-Altayó et al., 2020) | | | |
| KCl-induced contraction (aorta) | Increased (Jiménez-Altayó et al., 2020) | | | |
| Cranofacial features | | | | |
| Nose | Smaller [females] (Segura-Puimedon et al., 2014) | | | |
| Mandible | Shorter (Segura-Puimedon et al., 2014) | | | |
| Skull (A/P axis) | | Not shorter | Shorter, but less so than in D/P mice (Li et al., 2009) | Shorter (Li et al., 2009) |
| Neuroanatomy | | | | |
| Gross anatomy | Normal (Segura-Puimedon et al., 2014) | | | |
| Brain weight | | Normal (Li et al., 2009) | Reduced (~10%) (Li et al., 2009) | Reduced (~14%) (Li et al., 2009) |
| Amygdala | Reduced cell count (Segura-Puimedon et al., 2014) | | | |
| Hippocampus | Reduced CA3 area (Segura- | | | |

| | | | | |
|---|---|---|---|---|
| | Puimedon et al., 2014) | | | |
| Cortical thickness | Normal (Dasilva et al., 2020) | Increased neural density in layer V (Li et al., 2009) | | Increased neural density in layer V (Li et al., 2009) |
| Lateral ventricle | | Reduced size (Li et al., 2009) | Normal (Li et al., 2009) | Reduced size (Li et al., 2009) |
| Coherence (K/X anaesthesia; frontal cortex) | Less gamma band (Dasilva et al., 2020) | | | |
| Non-social behaviour | | | | |
| Wire test | Impaired (Segura-Puimedon et al., 2014) | | | Impaired (2) |
| Grip strength | | | | |
| Single pellet food retrieval (fine motricity) | | | | |
| Hot plate test | | | | |
| Gait pattern | Normal (Segura-Puimedon et al., 2014)<br>Slightly abnormal (Rahn et al., 2021) | | | |
| Open field (distance) | Normal (Segura-Puimedon et al., 2014; Kopp et al., 2019) | Normal (Li et al., 2009) | Decreased (Li et al., 2009) | Decreased (Li et al., 2009) |
| Open field (anxiety) | Normal (Segura-Puimedon et al., 2014; Kopp et al., 2019) | Normal (Li et al., 2009) | Normal (Li et al., 2009) | More anxious (Li et al., 2009) |
| Elevated maze | | | | |
| Rotarod | Impaired (Segura-Puimedon et al., 2014; Ortiz-Romero et al., 2021);<br>Not impaired (Kopp et al., 2019) | Impaired (Li et al., 2009) | Slightly impaired (Li et al., 2009) | Impaired (Li et al., 2009) |
| Acoustic startle | Increased (Segura- | | | |

| | | | |
|---|---|---|---|
| | Puimedon et al., 2014) | | |
| Prepulse inhibition | Normal (Kopp et al., 2019) | Reduced inhibition (Li et al., 2009) | Normal (Li et al., 2009) |
| Fear conditioning | Decreased (Segura-Puimedon et al., 2014; Nygaard et al., 2022) Increased (Kopp et al., 2019) | | Decreased (Li et al., 2009)) |
| Visual memory | Normal (Segura-Puimedon et al., 2014) | | |
| Water maze (distance) | Reduced (Segura-Puimedon et al., 2014) | | |
| Water maze (speed) | Reduced (Segura-Puimedon et al., 2014) | | |
| Water maze (probe) | Improved (p = 0.07) (Segura-Puimedon et al., 2014) | | |
| Marble burying | Less marbles buried (Kopp et al., 2019; Ortiz-Romero et al., 2021) | | |
| Ledge test | Impaired balance (Kopp et al., 2019) | | |
| Elevated plus maze | Normal (Kopp et al., 2019) | | |
| Spontaneous alternation test (spatial working memory) | Impaired (Borralleras et al., 2016) | | |
| Social behaviour | | | |
| Social approach | Increased (Segura-Puimedon et al., 2014; Ortiz- | | |

| | | | | |
|---|---|---|---|---|
| | Romero et al., 2021) | | | |
| Dyadic social interaction | | | | |
| Three-chamber test | Increased sociability (Kopp et al., 2019) | Increased sociability (Li et al., 2009) | Normal (Li et al., 2009) | Increased sociability (Li et al., 2009) |
| Tube test | Submissive (Kopp et al., 2019) | Submissive (Li et al., 2009) | Normal (Li et al., 2009) | Submissive (Li et al., 2009) |
| USV (maternal separation) | Less calls (Kopp et al., 2019; Giannoccaro et al., 2023) | | | |
| Electrophysiology | | | | |
| Hippocampal LTP | Impaired (Borralleras et al., 2016) | | | |

**Table S2. Statistical analysis of Fig. S12**

The statistical tests (repeated measures ANOVA) for the peak ratios were done after root-transformation of the ratios.

| | | Between effects | df | F | P | | Within effects | df | F | P |
|---|---|---|---|---|---|---|---|---|---|---|
| ABR threshold | | Intercept | 1 | 2288 | 0 | | Frequency | 4 | 378 | 0 |
| | | Genotype | 1 | 0.02 | 0.89 | | Genotype:Frequency | 4 | 0.44 | 0.78 |
| | | Error | 23 | | | | Error | 92 | | |
| 12 kHz | Peak I | Intercept | 1 | 733 | 0 | | Intensity | 11 | 128 | 0 |
| | | Genotype | 1 | 1.91 | 0.18 | | Genotype:Intensity | 11 | 1.06 | 0.39 |
| | | Error | 23 | | | | Error | 253 | | |
| | Peak IV | Intercept | 1 | 766 | 0 | | Intensity | 11 | 66 | 0 |
| | | Genotype | 1 | 3.89 | 0.06 | | Genotype:Intensity | 11 | 6.3 | 3 $10^{-9}$ |
| | | Error | 23 | | | | Error | 253 | | |
| | Peak VI | Intercept | 1 | 549 | 0 | | Intensity | 11 | 29 | 0 |
| | | Genotype | 1 | 0.79 | 0.38 | | Genotype:Intensity | 11 | 1.3 | 0.23 |
| | | Error | 23 | | | | Error | 253 | | |
| | Ratio IV/I | Intercept | 1 | 4768 | 0 | | Intensity | 8 | 29 | 0 |
| | | Genotype | 1 | 0.66 | 0.42 | | Genotype:Intensity | 8 | 4.6 | 4 $10^{-5}$ |
| | | Error | 23 | | | | Error | 184 | | |
| | Ratio VI/I | Intercept | 1 | 1373 | 0 | | Intensity | 8 | 2.2 | 0.03 |
| | | Genotype | 1 | 5.8 | 0.02 | | Genotype:Intensity | 8 | 0.9 | 0.5 |
| | | Error | 23 | | | | Error | 184 | | |
| | Ratio VI/IV | Intercept | 1 | 1699 | 0 | | Intensity | 8 | 2.3 | 0.02 |
| | | Genotype | 1 | 9.8 | 0.005 | | Genotype:Intensity | 8 | 0.6 | 0.7 |
| | | Error | 23 | | | | Error | 184 | | |
| 16 kHz | Peak I | Intercept | 1 | 351 | 2 $10^{-5}$ | | Intensity | 9 | 126 | 0 |
| | | Genotype | 1 | 0.17 | 0.69 | | Genotype:Intensity | 9 | 0.95 | 0.48 |
| | | Error | 23 | | | | Error | 207 | | |
| | Peak IV | Intercept | 1 | 593 | 0 | | Intensity | 9 | 57 | 0 |
| | | Genotype | 1 | 3.4 | 0.077 | | Genotype:Intensity | 9 | 2.2 | 0.023 |
| | | Error | 23 | | | | Error | 207 | | |
| | Peak VI | Intercept | 1 | 405 | 0 | | Intensity | 9 | 53 | 0 |
| | | Genotype | 1 | 6.9 | 0.015 | | Genotype:Intensity | 9 | 2.7 | 0.0057 |
| | | Error | 23 | | | | Error | 207 | | |
| | Ratio IV/I | Intercept | 1 | 3284 | 0 | | Intensity | 8 | 60 | 0 |
| | | Genotype | 1 | 5.3 | 0.030 | | Genotype:Intensity | 8 | 1.8 | 0.075 |
| | | Error | 23 | | | | Error | 184 | | |
| | Ratio VI/I | Intercept | 1 | 1889 | 0 | | Intensity | 8 | 1 | 0.67 |
| | | Genotype | 1 | 3.9 | 0.060 | | Genotype:Intensity | 8 | 0.5 | 0.86 |
| | | Error | 23 | | | | Error | 184 | | |
| | Ratio VI/IV | Intercept | 1 | 1329 | 0 | | Intensity | 8 | 10 | 6 $10^{-12}$ |
| | | Genotype | 1 | 13.6 | 0.0012 | | Genotype:Intensity | 8 | 2.1 | 0.039 |
| | | Error | 23 | | | | Error | 184 | | |

**Table S3. Group averages, quantiles and post-hoc testing of within-interaction terms of Fig. S12**

Differences are indicated and tested for significance if the main effect of genotype or the genotype within-interaction term or both were significant (Table S2). P-values were corrected following Tukey-Kramer. The statistical tests for the ratios were done after root-transformation of the ratios. The reported mean and quartiles for the ratios are not transformed.

| | | WT | | LD | | Tests of within-interaction terms | | |
|---|---|---|---|---|---|---|---|---|
| | | Mean | Quartiles | Mean | Quartiles | Genotype:Frequency | | |
| | Frequency (kHz) | Hearing thresholds (dB SPL) | | | | difference | SE | P |
| Hearing thresholds | 4 | 43.6 | [40.0, 45.0] | 43.6 | [40.0, 45.0] | | | |
| | 8 | 18.6 | [15.0, 20.0] | 20.0 | [20.0, 20.0] | | | |
| | 12 | 10.7 | [10.0, 13.8] | 9.5 | [10.0, 10.0] | | | |
| | 16 | 13.2 | [10.0, 15.0] | 13.2 | [10.0, 15.0] | | | |
| | 32 | 39.6 | [35.0, 43.8] | 38.6 | [35.0, 42.5] | | | |
| | | | | | | Genotype:Intensity | | |
| | Intensity (dB SPL) | Amplitude (µV) | | | | difference | SE | P |
| 12 kHz – Peak I | 15 | 0.08 | [0.05, 0.10] | 0.08 | [0.06, 0.10] | | | |
| | 20 | 0.12 | [0.10, 0.14] | 0.11 | [0.09, 0.13] | | | |
| | 25 | 0.17 | [0.14, 0.18] | 0.15 | [0.14, 0.17] | | | |
| | 30 | 0.20 | [0.18, 0.22] | 0.18 | [0.16, 0.21] | | | |
| | 35 | 0.25 | [0.22, 0.27] | 0.22 | [0.20, 0.24] | | | |
| | 40 | 0.29 | [0.24, 0.30] | 0.25 | [0.23, 0.29] | | | |
| | 45 | 0.30 | [0.27, 0.31] | 0.26 | [0.23, 0.31] | | | |
| | 50 | 0.31 | [0.27, 0.33] | 0.29 | [0.28, 0.33] | | | |
| | 55 | 0.33 | [0.25, 0.36] | 0.32 | [0.29, 0.38] | | | |
| | 60 | 0.40 | [0.28, 0.38] | 0.38 | [0.32, 0.42] | | | |
| | 70 | 0.54 | [0.43, 0.59] | 0.49 | [0.46, 0.54] | | | |

| | | | | | | | | |
|---|---|---|---|---|---|---|---|---|
| | 80 | 0.87 | [0.77, 0.89] | 0.73 | [0.65, 0.79] | | | |
| 12 kHz – Peak IV | 15 | 0.11 | [0.08, 0.13] | 0.14 | [0.11, 0.17] | 0.03 | 0.02 | 0.14 |
| | 20 | 0.18 | [0.14, 0.23] | 0.19 | [0.15, 0.23] | 0.01 | 0.02 | 0.68 |
| | 25 | 0.24 | [0.20, 0.29] | 0.23 | [0.19, 0.26] | -0.01 | 0.02 | 0.79 |
| | 30 | 0.27 | [0.25, 0.31] | 0.27 | [0.25, 0.31] | 0.00 | 0.02 | 0.92 |
| | 35 | 0.30 | [0.28, 0.32] | 0.30 | [0.28, 0.34] | 0.00 | 0.02 | 0.93 |
| | 40 | 0.35 | [0.33, 0.37] | 0.34 | [0.31, 0.40] | -0.01 | 0.03 | 0.77 |
| | 45 | 0.37 | [0.33, 0.40] | 0.36 | [0.32, 0.40] | -0.01 | 0.02 | 0.57 |
| | 50 | 0.41 | [0.37, 0.42] | 0.36 | [0.31, 0.43] | -0.05 | 0.03 | 0.085 |
| | 55 | 0.43 | [0.40, 0.44] | 0.38 | [0.33, 0.43] | -0.05 | 0.03 | 0.077 |
| | 60 | 0.48 | [0.39, 0.48] | 0.39 | [0.34, 0.43] | -0.09 | 0.05 | 0.099 |
| | 70 | 0.56 | [0.44, 0.59] | 0.40 | [0.33, 0.46] | -0.16 | 0.06 | 0.015 |
| | 80 | 0.73 | [0.53, 0.88] | 0.48 | [0.38, 0.58] | -0.25 | 0.09 | 0.0086 |
| 12 kHz – Peak VI | 15 | 0.07 | [0.03, 0.11] | 0.06 | [0.04, 0.08] | | | |
| | 20 | 0.09 | [0.06, 0.12] | 0.06 | [0.03, 0.08] | | | |
| | 25 | 0.08 | [0.06, 0.10] | 0.07 | [0.05, 0.09] | | | |
| | 30 | 0.09 | [0.04, 0.11] | 0.12 | [0.10, 0.14] | | | |
| | 35 | 0.11 | [0.07, 0.13] | 0.11 | [0.08, 0.15] | | | |
| | 40 | 0.12 | [0.07, 0.14] | 0.13 | [0.09, 0.15] | | | |
| | 45 | 0.12 | [0.07, 0.16] | 0.15 | [0.11, 0.21] | | | |
| | 50 | 0.12 | [0.10, 0.14] | 0.15 | [0.11, 0.19] | | | |
| | 55 | 0.14 | [0.10, 0.16] | 0.17 | [0.15, 0.21] | | | |
| | 60 | 0.15 | [0.11, 0.19] | 0.20 | [0.14, 0.23] | | | |

| | | | | | | | | |
|---|---|---|---|---|---|---|---|---|
| | 70 | 0.20 | [0.16, 0.23] | 0.17 | [0.14, 0.19] | | | |
| | 80 | 0.28 | [0.23, 0.31] | 0.29 | [0.24, 0.33] | | | |
| | | **Ratio** | | | | | | |
| **12 kHz – Ratio IV/I** | 30 | 1.35 | [1.22, 1.51] | 1.49 | [1.39, 1.64] | 0.06 | 0.05 | 0.18 |
| | 35 | 1.26 | [1.14, 1.33] | 1.37 | [1.22, 1.55] | 0.05 | 0.05 | 0.28 |
| | 40 | 1.24 | [1.14, 1.33] | 1.37 | [1.22, 1.48] | 0.06 | 0.04 | 0.18 |
| | 45 | 1.27 | [1.16, 1.45] | 1.43 | [1.19, 1.56] | 0.07 | 0.05 | 0.18 |
| | 50 | 1.37 | [1.26, 1.47] | 1.28 | [1.14, 1.42] | -0.04 | 0.04 | 0.39 |
| | 55 | 1.38 | [1.21, 1.50] | 1.19 | [1.03, 1.36] | -0.08 | 0.05 | 0.11 |
| | 60 | 1.31 | [1.17, 1.49] | 1.07 | [0.86, 1.29] | -0.11 | 0.06 | 0.070 |
| | 70 | 1.09 | [0.86, 1.23] | 0.81 | [0.69, 0.95] | -0.14 | 0.05 | 0.013 |
| | 80 | 0.86 | [0.63, 1.12] | 0.66 | [0.52, 0.87] | -0.11 | 0.06 | 0.077 |
| **12 kHz – Ratio VI/I** | 30 | 0.46 | [0.21, 0.49] | 0.64 | [0.51, 0.73] | 0.16 | 0.08 | 0.052 |
| | 35 | 0.44 | [0.27, 0.52] | 0.50 | [0.37, 0.61] | 0.06 | 0.06 | 0.37 |
| | 40 | 0.42 | [0.24, 0.46] | 0.52 | [0.34, 0.67] | 0.08 | 0.07 | 0.24 |
| | 45 | 0.41 | [0.24, 0.55] | 0.64 | [0.41, 0.71] | 0.15 | 0.08 | 0.057 |
| | 50 | 0.39 | [0.30, 0.48] | 0.53 | [0.40, 0.66] | 0.10 | 0.06 | 0.12 |
| | 55 | 0.44 | [0.24, 0.51] | 0.54 | [0.41, 0.68] | 0.10 | 0.07 | 0.19 |
| | 60 | 0.42 | [0.26, 0.61] | 0.54 | [0.38, 0.65] | 0.09 | 0.06 | 0.16 |
| | 70 | 0.38 | [0.26, 0.42] | 0.36 | [0.29, 0.40] | -0.02 | 0.05 | 0.70 |
| | 80 | 0.32 | [0.27, 0.37] | 0.39 | [0.35, 0.46] | 0.06 | 0.03 | 0.098 |
| **12 kHz – Ratio VI/IV** | 30 | 0.35 | [0.15, 0.37] | 0.43 | [0.36, 0.47] | 0.10 | 0.07 | 0.15 |
| | 35 | 0.35 | [0.24, 0.47] | 0.38 | [0.25, 0.51] | 0.03 | 0.06 | 0.65 |
| | 40 | 0.34 | [0.19, 0.39] | 0.39 | [0.27, 0.51] | 0.04 | 0.06 | 0.48 |

| | | | | | | | | |
|---|---|---|---|---|---|---|---|---|
| | 45 | 0.34 | [0.19, 0.42] | 0.43 | [0.30, 0.54] | 0.19 | 0.06 | 0.16 |
| | 50 | 0.29 | [0.22, 0.36] | 0.41 | [0.31, 0.53] | 0.10 | 0.05 | 0.058 |
| | 55 | 0.33 | [0.19, 0.40] | 0.46 | [0.39, 0.54] | 0.12 | 0.06 | 0.050 |
| | 60 | 0.33 | [0.26, 0.39] | 0.51 | [0.38, 0.57] | 0.14 | 0.05 | 0.015 |
| | 70 | 0.37 | [0.25, 0.50] | 0.45 | [0.33, 0.55] | 0.07 | 0.05 | 0.21 |
| | 80 | 0.42 | [0.25, 0.54] | 0.66 | [0.43, 0.78] | 0.16 | 0.07 | 0.033 |
| | | **Amplitude (μV)** | | | | | | |
| **16 kHz – Peak I** | 25 | 0.10 | [0.09, 0.11] | 0.12 | [0.09, 0.15] | | | |
| | 30 | 0.15 | [0.12, 0.16] | 0.15 | [0.12, 0.19] | | | |
| | 35 | 0.20 | [0.17, 0.23] | 0.18 | [0.13, 0.23] | | | |
| | 40 | 0.24 | [0.20, 0.28] | 0.28 | [0.20, 0.31] | | | |
| | 45 | 0.27 | [0.23, 0.30] | 0.27 | [0.23, 0.31] | | | |
| | 50 | 0.30 | [0.24, 0.34] | 0.30 | [0.24, 0.33] | | | |
| | 55 | 0.34 | [0.28, 0.38] | 0.34 | [0.28, 0.37] | | | |
| | 60 | 0.45 | [0.34, 0.48] | 0.54 | [0.34, 0.72] | | | |
| | 70 | 0.56 | [0.46, 0.59] | 0.53 | [0.46, 0.59] | | | |
| | 80 | 0.78 | [0.67, 0.91] | 0.82 | [0.64, 0.99] | | | |
| **16 kHz – Peak IV** | 25 | 0.24 | [0.21, 0.28] | 0.19 | [0.14, 0.24] | -0.05 | 0.03 | 0.098 |
| | 30 | 0.28 | [0.23, 0.31] | 0.26 | [0.26, 0.30] | -0.01 | 0.03 | 0.61 |
| | 35 | 0.33 | [0.30, 0.35] | 0.31 | [0.28, 0.35] | -0.01 | 0.02 | 0.62 |
| | 40 | 0.40 | [0.36, 0.43] | 0.37 | [0.29, 0.44] | -0.03 | 0.04 | 0.54 |
| | 45 | 0.43 | [0.38, 0.48] | 0.36 | [0.31, 0.44] | -0.07 | 0.04 | 0.079 |
| | 50 | 0.47 | [0.41, 0.52] | 0.40 | [0.31, 0.49] | -0.07 | 0.04 | 0.087 |
| | 55 | 0.50 | [0.46, 0.55] | 0.44 | [0.38, 0.51] | -0.06 | 0.04 | 0.12 |

| | | | | | | | | |
|---|---|---|---|---|---|---|---|---|
| | 60 | 0.54 | [0.47, 0.62] | 0.50 | [0.37, 0.55] | -0.04 | 0.06 | 0.47 |
| | 70 | 0.54 | [0.42, 0.64] | 0.42 | [0.39, 0.47] | -0.13 | 0.05 | 0.011 |
| | 80 | 0.62 | [0.51, 0.70] | 0.48 | [0.36, 0.64] | -0.14 | 0.07 | 0.048 |
| **16 kHz – Peak VI** | 25 | 0.04 | [0.01, 0.05] | 0.06 | [0.04, 0.07] | 0.02 | 0.01 | 0.17 |
| | 30 | 0.05 | [0.03, 0.05] | 0.08 | [0.04, 0.12] | 0.04 | 0.02 | 0.078 |
| | 35 | 0.08 | [0.07, 0.09] | 0.08 | [0.05, 0.12] | 0.00 | 0.02 | 0.81 |
| | 40 | 0.09 | [0.07, 0.12] | 0.11 | [0.08, 0.14] | 0.01 | 0.01 | 0.38 |
| | 45 | 0.09 | [0.08, 0.13] | 0.13 | [0.11, 0.15] | 0.04 | 0.02 | 0.089 |
| | 50 | 0.10 | [0.09, 0.11] | 0.13 | [0.10, 0.15] | 0.03 | 0.02 | 0.18 |
| | 55 | 0.12 | [0.09, 0.15] | 0.13 | [0.09, 0.15] | 0.01 | 0.02 | 0.76 |
| | 60 | 0.13 | [0.10, 0.16] | 0.18 | [0.14, 0.21] | 0.05 | 0.02 | 0.021 |
| | 70 | 0.16 | [0.11, 0.20] | 0.19 | [0.15, 0.22] | 0.03 | 0.03 | 0.30 |
| | 80 | 0.23 | [0.19, 0.23] | 0.34 | [0.27, 0.40] | 0.11 | 0.04 | 0.0058 |
| | | **Ratio** | | | | | | |
| **16 kHz – Ratio IV/I** | 30 | 2.05 | [1.58, 2.19] | 1.88 | [1.68, 2.03] | -0.05 | 0.08 | 0.56 |
| | 35 | 1.72 | [1.50, 1.87] | 1.91 | [1.52, 2.14] | 0.06 | 0.07 | 0.43 |
| | 40 | 1.69 | [1.51, 1.83] | 1.40 | [1.11, 1.61] | -0.13 | 0.05 | 0.026 |
| | 45 | 1.66 | [1.45, 1.70] | 1.39 | [1.19, 1.66] | -0.11 | 0.06 | 0.077 |
| | 50 | 1.67 | [1.41, 1.88] | 1.37 | [1.20, 1.57] | -0.12 | 0.06 | 0.059 |
| | 55 | 1.57 | [1.41, 1.63] | 1.32 | [1.11, 1.56] | -0.11 | 0.06 | 0.075 |
| | 60 | 1.27 | [1.08, 1.38] | 0.99 | [0.86, 1.05] | -0.13 | 0.05 | 0.0095 |
| | 70 | 1.01 | [0.86, 1.11] | 0.82 | [0.60, 0.98] | -0.10 | 0.05 | 0.034 |
| | 80 | 0.83 | [0.68, 0.96] | 0.58 | [0.45, 0.69] | -0.15 | 0.05 | 0.0058 |
| **16 kH** | 30 | 0.37 | [0.20, 0.32] | 0.51 | [0.25, 0.85] | | | |

| | | | | | | | | |
|---|---|---|---|---|---|---|---|---|
| | 35 | 0.40 | [0.31, 0.49] | 0.49 | [0.32, 0.57] | | | |
| | 40 | 0.40 | [0.31, 0.50] | 0.42 | [0.26, 0.52] | | | |
| | 45 | 0.38 | [0.22, 0.55] | 0.51 | [0.40, 0.65] | | | |
| | 50 | 0.39 | [0.24, 0.56] | 0.44 | [0.34, 0.53] | | | |
| | 55 | 0.38 | [0.30, 0.45] | 0.37 | [0.26, 0.48] | | | |
| | 60 | 0.31 | [0.21, 0.41] | 0.37 | [0.29, 0.46] | | | |
| | 70 | 0.29 | [0.22, 0.35] | 0.36 | [0.29, 0.42] | | | |
| | 80 | 0.30 | [0.23, 0.33] | 0.43 | [0.34, 0.51] | | | |
| **16 kHz – Ratio VI/IV** | 30 | 0.19 | [0.10, 0.20] | 0.31 | [0.13, 0.52] | 0.09 | 0.09 | 0.32 |
| | 35 | 0.24 | [0.19, 0.28] | 0.26 | [0.16, 0.35] | 0.01 | 0.05 | 0.90 |
| | 40 | 0.24 | [0.21, 0.30] | 0.32 | [0.21, 0.42] | 0.06 | 0.04 | 0.15 |
| | 45 | 0.22 | [0.16, 0.29] | 0.38 | [0.26, 0.46] | 0.16 | 0.07 | 0.039 |
| | 50 | 0.23 | [0.17, 0.25] | 0.35 | [0.20, 0.44] | 0.10 | 0.05 | 0.065 |
| | 55 | 0.25 | [0.20, 0.28] | 0.30 | [0.19, 0.42] | 0.04 | 0.04 | 0.41 |
| | 60 | 0.24 | [0.19, 0.31] | 0.38 | [0.27, 0.50] | 0.12 | 0.04 | 0.0040 |
| | 70 | 0.31 | [0.21, 0.41] | 0.47 | [0.34, 0.56] | 0.14 | 0.05 | 0.020 |
| | 80 | 0.39 | [0.31, 0.47] | 0.83 | [0.54, 1.11] | 0.28 | 0.07 | 0.00064 |
